# Plasticity in gustatory and nociceptive neurons controls decision making in *C. elegans* salt navigation

**DOI:** 10.1101/2021.03.19.436114

**Authors:** Martijn P.J. Dekkers, Felix Salfelder, Tom Sanders, Oluwatoroti Umuerri, Netta Cohen, Gert Jansen

**Affiliations:** Department of Cell Biology, Erasmus University Medical Centre, Rotterdam, The Netherlands; School of Computing, University of Leeds, Leeds, United Kingdom; REHAB Basel, Basel, Switzerland

## Abstract

A conventional understanding of perception assigns sensory organs the role of capturing the environment. Better sensors result in more accurate encoding of stimuli, allowing for cognitive processing downstream. Here we show that plasticity in sensory neurons mediates a behavioral switch in *C. elegans* between attraction to NaCl in naïve animals and avoidance of NaCl in preconditioned animals, called gustatory plasticity. Ca^2+^ imaging in ASE and ASH NaCl sensing neurons reveals multiple cell-autonomous and distributed circuit adaptation mechanisms. A computational model quantitatively accounts for observed behaviors and reveals roles for sensory neurons in the control and modulation of motor behaviors, decision making and navigational strategy. Sensory adaptation dynamically alters the encoding of the environment. Rather than encoding the stimulus directly, therefore, we propose that these *C. elegans* sensors dynamically encode a context-dependent value of the stimulus. Our results demonstrate how adaptive sensory computation can directly control an animal’s behavioral state.

## Introduction

Decision making refers to the process of choosing among distinct actions as a function of the estimated value of their consequences. In expected utility theory, rational agents assign a subjective value (or expected utility) to a particular action^1^. The subjective value of different outcomes may be context dependent (e.g., the perceived value of food may be hunger-dependent, as is the cost of food deprivation^2,3^) and limited by noise or partial information^1^. In fact, in all but the simplest behaviors, the information available to an individual will not directly determine the actual utility of different choices to the individual, implying that in the real world, people and animals need to infer, learn and dynamically adapt the estimated value of different actions. Thus, adaptation is a universal defining feature of animal behavior^4^.

Adaptive behavior refers to the ability of animals to change their actions in response to changes in the environment or in their internal state. Here, we study a form of short-term sensory adaptation, on timescales of 15 minutes or less, called gustatory plasticity^2,5,6^. Its defining feature in the nematode *Caenorhabditis elegans* is the dynamic balance of salt (NaCl) attraction and avoidance as a function of experience. Often a goal is implicit as the driver for adaptive behavior. In fact, as NaCl has little or no objective value to the animal, one might surmise that the expected reward of seeking salt is food. However, information about salt does not directly or reliably predict the presence of food. Gustatory plasticity may therefore be an adaptive mechanism for modulating the expected utility of following (or avoiding) salt concentration gradients, in search of food.

The response of *C. elegans* to NaCl is associated with food. Naïve animals, cultured in the presence of food and NaCl, will move up NaCl concentration gradients in search of bacteria^2,7,8^. Preconditioned animals, exposed for 15 minutes to 100 mM NaCl in the absence of food, avoid any NaCl concentration. This switch from attractive to aversive NaCl behavior is called gustatory plasticity^2,5,6^. Gustatory plasticity is reversible, lasting less than 5 minutes^6^. 30 minutes or longer exposure to NaCl in the absence of food induces stronger avoidance responses that rely on mostly independent mechanisms^5,9-13^.

While adaptive decision making typically involves an integration of sensory stimuli with information about prior experience and internal state, the neural basis for adaptive decision making is still far from understood. Most studies of adaptive behavior have focused on sites of multi-sensory and internal-state integration in downstream neurons and circuits^14^. We present evidence for the history-dependent modulation of value encoding and decision making by sensory neurons during gustatory plasticity in the nematode *C. elegans*.

Naïve *C. elegans* are attracted to NaCl concentrations of up to 200 mM but avoid higher NaCl concentrations^2,7,8^. Attraction to NaCl is primarily mediated by the bilaterally asymmetric ASE sensory neurons^15,16^. The left ASE neuron (ASEL) produces Ca^2+^ transients in response to increases in NaCl concentration^17^; the right neuron (ASER) responds to decreases in NaCl concentration^17^. Avoidance of dangerously high NaCl concentrations is mediated by the ASH neurons^7,18^. Previous studies have shown that the ASE and the ASH neurons are involved in in gustatory plasticity^2^. In addition, many signaling proteins involved have been identified, including serotonin, dopamine, glutamate and neuropeptide neurotransmission^2,5,6,9^. In this paper, we study adaptive mechanisms in ASEL, ASER and ASH to identify possible neuronal and circuit mechanisms of gustatory plasticity and to link neuronal dynamics with the animal’s adaptive behavior.

Cell specific Ca^2+^ imaging in awake animals identified three distinct forms of adaptation that occur in the absence of food and overlap with the timescale of gustatory plasticity: ASEL desensitization upon exposure to NaCl; ASER sensitization to NaCl; and ASH sensitization to considerably lower (non-toxic) levels of NaCl. An additional, fast form of dynamic-range adaptation is identified in ASE sensory neurons, resulting in a logarithmic response amplitude to changes of NaCl concentration, analogous to the Weber-Fechner law of sensory perception^19,20^. Using computational models, we identify a hierarchy of molecular, cellular and distributed circuit mechanisms that capture our Ca^2+^ imaging results in sensory neurons. Simulations of model animals in a virtual assay environment captured the behavioral switch from attraction to avoidance in gustatory plasticity. Our experimental results and computational model point to a number of predictions: First, ASH sensitization is necessary and sufficient to explain the behavioral switch in gustatory plasticity. Second, bilateral asymmetries in ASE adaptation limit the animals’ ability to follow NaCl gradients but make these neurons excellent adaptive encoders of context-and history-dependent value that drives different motor actions on different timescales. Finally, we postulate a novel role of sensory adaptation in setting the balance of exploration and exploitation in ecologically relevant scenarios and use our computational framework to support this conjecture in a simplified virtual assay.

## Results

### Naïve sensory responses to NaCl

What drives the behavioral switch between NaCl attraction and avoidance during gustatory plasticity? Before addressing this question, we determined the range of the naïve responses to NaCl of the ASEL, ASER, and ASH neurons, using the Ca^2+^ reporter Yellow Cameleon^21,22^. Similar to previous findings^17^ ASEL neurons produced Ca^2+^ transients in response to a 3 seconds exposure to both low and high NaCl concentrations, with strongest responses to 200 mM NaCl (**Fig. 1a,b**; **Supplementary Table 1**). ASH neurons are known to yield Ca^2+^ transients in response to osmotic stimuli^23^. We recorded Ca^2+^ transients in ASH neurons in response to a 3 seconds exposure to various NaCl concentrations. We found a gradual increase in the fraction of animals that responded (depicted as the response index, RI, **Supplementary Table 1)** and in the amplitude of Ca^2+^ transients with increasing concentrations of NaCl, resulting in strong Ca^2+^ fluxes in response to 300 mM and 500 mM NaCl, but only a small fraction of animals responded to 100 or 200 mM NaCl and the associated Ca^2+^ fluxes were weak (**Fig. 1d,e**; **Supplementary Table 1**).

**Figure 1.**
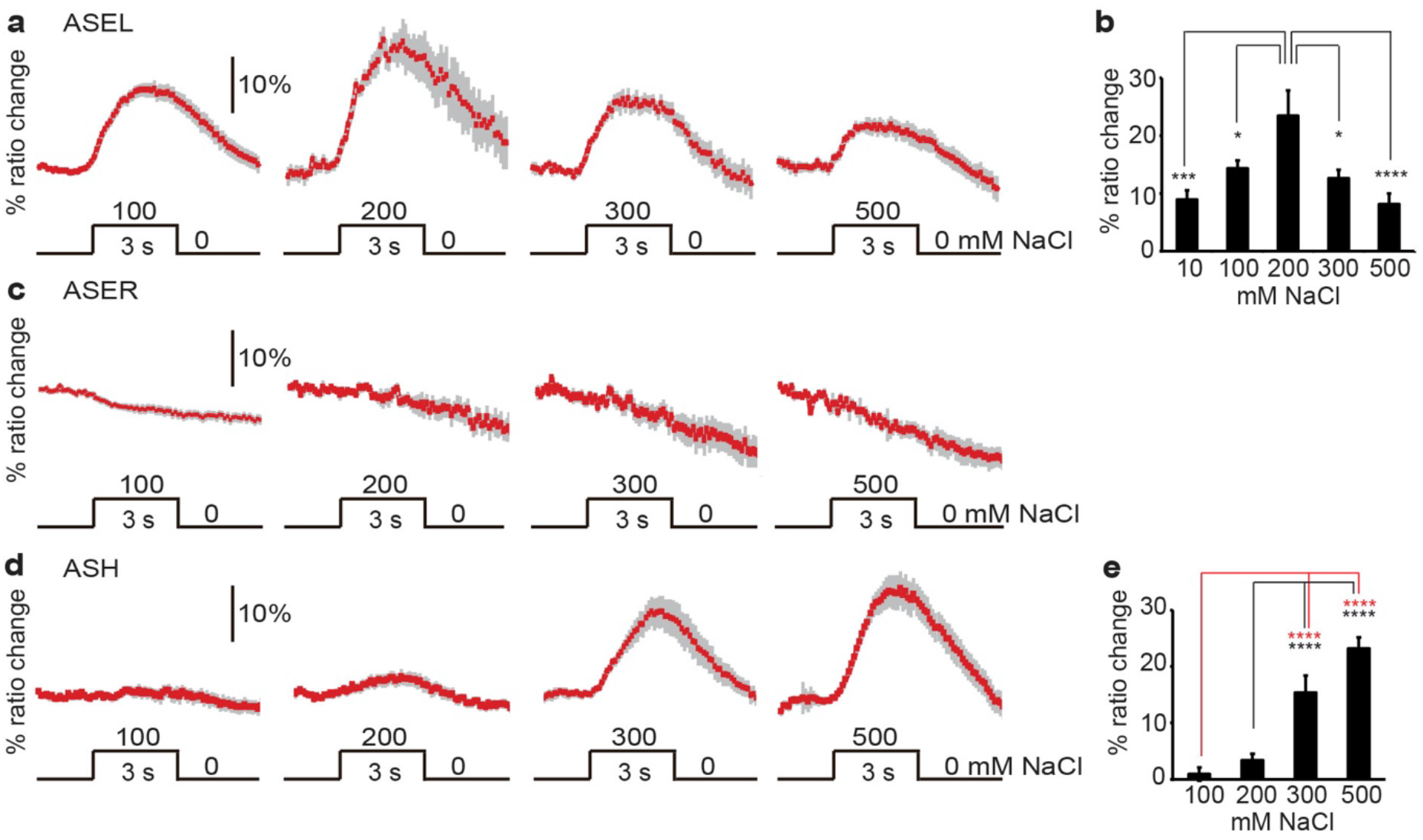
Ca^2+^ responses of ASEL, ASER and ASH neurons to brief NaCl exposure. Animals were exposed for 3 seconds to different concentrations of NaCl from a baseline of 0 mM. (**a**) Average Ca^2+^ transients (± SEM in gray) in ASEL in response to 100 −500 mM NaCl; 100 mM: n = 37, 200 mM: n = 9, 300 mM: n = 15, 500 mM: n = 8. (**b**) Average maximum ratio changes (± SEM) in ASEL: responses to 10 (n=10), 100 (n=37), 300 (n=15) and 500 (n=8) mM were significantly different from the response to 200 mM. (**c**) Average Ca^2+^ transients (± SEM in gray) in ASER in response to 100 −500 mM NaCl; 100 mM: n = 30, 200 mM: n = 5, 300 mM: n = 9, 500 mM: n = 4. No statistically significant differences were observed (p>0.05). (d) Average Ca^2+^ transients (± SEM) in ASH after exposure to 100 −500 mM NaCl; 100 mM: n = 18, 200 mM: n = 18, 300 mM: n = 22, 500 mM: n = 19. (e) Average maximum ratio changes (± SEM) in ASH: responses to 300 and 500 mM were significantly different from the responses to 100 and 200 mM. Traces indicate average percentage change in R/R_o_ where R is the fluorescence emission ratio and R_o_ the baseline fluorescence emission ratio before exposure to NaCl. Statistical significance *: p<0.05, *** p<0.005, ****: p<0.001 (non-significant differences, p>0.05, have not been indicated).

### Prolonged exposure to NaCl sensitizes ASER

In contrast to previous studies which have found ASER responses to decreases in NaCl concentrations^17,24,25^, we did not find responses to NaCl concentration decrease after a 3 seconds exposure, either at low or at high concentrations (**Fig. 1c**; **Supplementary Table 1**). This surprising result, combined with the fact that the ASER neuron is known to contribute to NaCl chemotaxis^17,24,25^, led us to conjecture that ASER responses may depend on its history of exposure to NaCl. To test this hypothesis, we measured ASER Ca^2+^ responses in animals exposed to 100 mM NaCl for 30 seconds to 10 minutes. We found that the fraction of animals that responded, as well as the amplitude of the response increased with exposure time (**Fig. 2a,b**; **Supplementary Table 1**), indicating that ASER is gradually sensitized by prolonged exposure to NaCl and confirming that its responses are consistent with positive (attractive) chemotaxis over behaviorally relevant timescales^25,26^.

**Figure 2.**
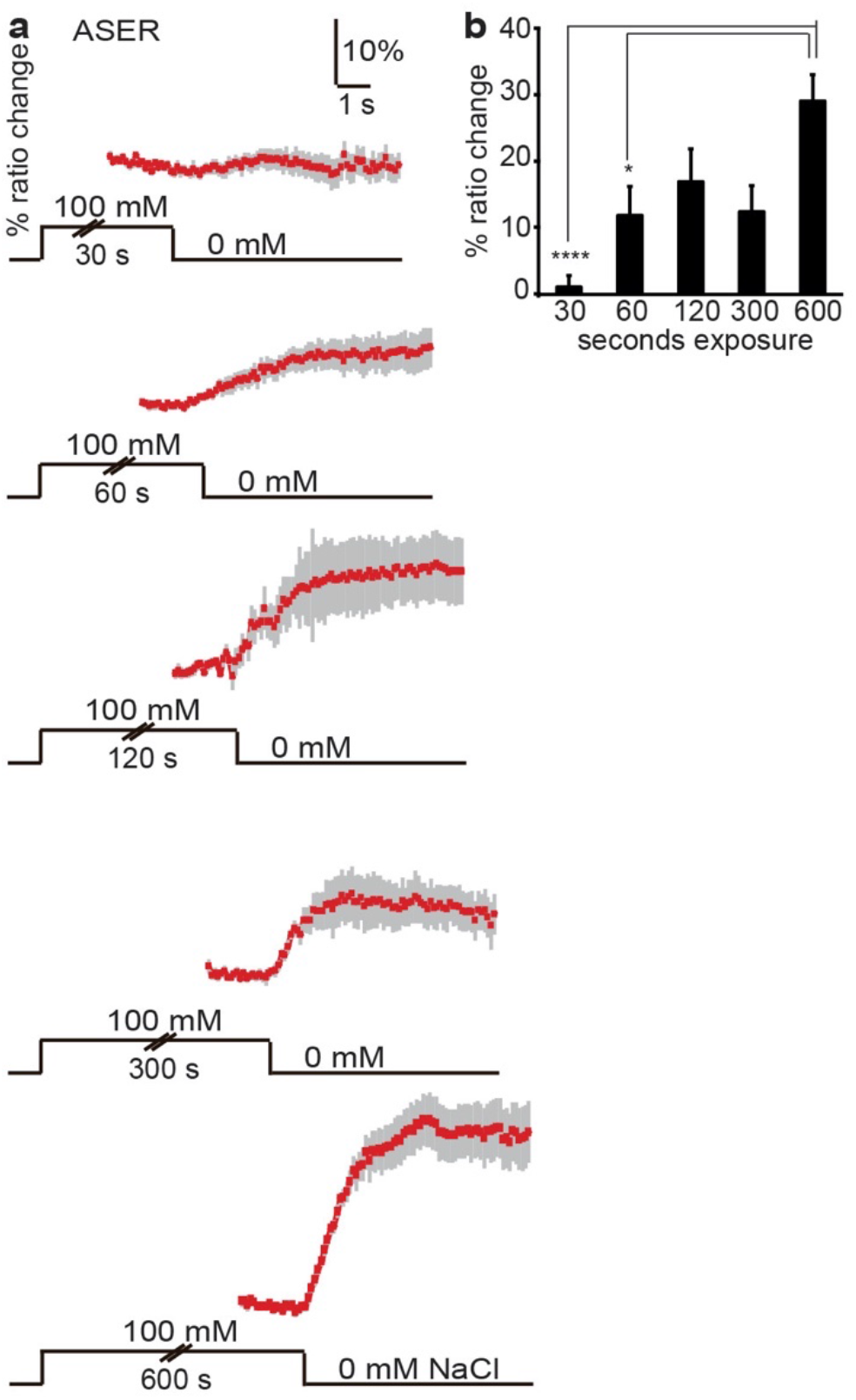
Prolonged exposure to NaCl sensitizes ASER. (**a**) Average Ca^2+^ transients (± SEM) in ASER in response to a decrease in NaCl concentration from 100 mM to 0 mM after 30 −600 seconds exposure. 30 seconds exposure to 100 mM NaCl did not result in a response in ASER, but longer exposures did. (**b**) Average maximum ratio changes (± SEM) in ASER after 30 −600 seconds exposure. 30 seconds: n = 8, 1 minute: n = 15, 2 minutes: n = 4, 5 minutes: n = 5, 10 minutes: n = 29. Statistical significance *: p<0.05, ****: p<0.001 (non-significant differences, p>0.05, have not been indicated).

### Prolonged exposure to NaCl desensitizes ASEL

To determine whether ASEL responses are also modulated by pre-exposure, we tested ASEL responses to 100 mM NaCl after a period of pre-exposure. Animals were pre-exposed to 100 mM NaCl for periods ranging from 60 seconds to 10 minutes, followed by a 60 seconds wash (**Fig. 3a**). Ca^2+^ responses in ASEL neurons were strongly reduced or even abolished after 5 or 10 minutes of pre-exposure to 100 mM NaCl, but unaffected after 1 or 2 minutes of pre-exposure (**Fig. 3a,b**; **Supplementary Table 1**). These results correlate well with behavioral assays that showed reduced attraction to NaCl with increasing pre-exposure times (**Supplementary Fig. 1**), as reported previously for the response to sodium acetate^6^.

**Figure 3.**
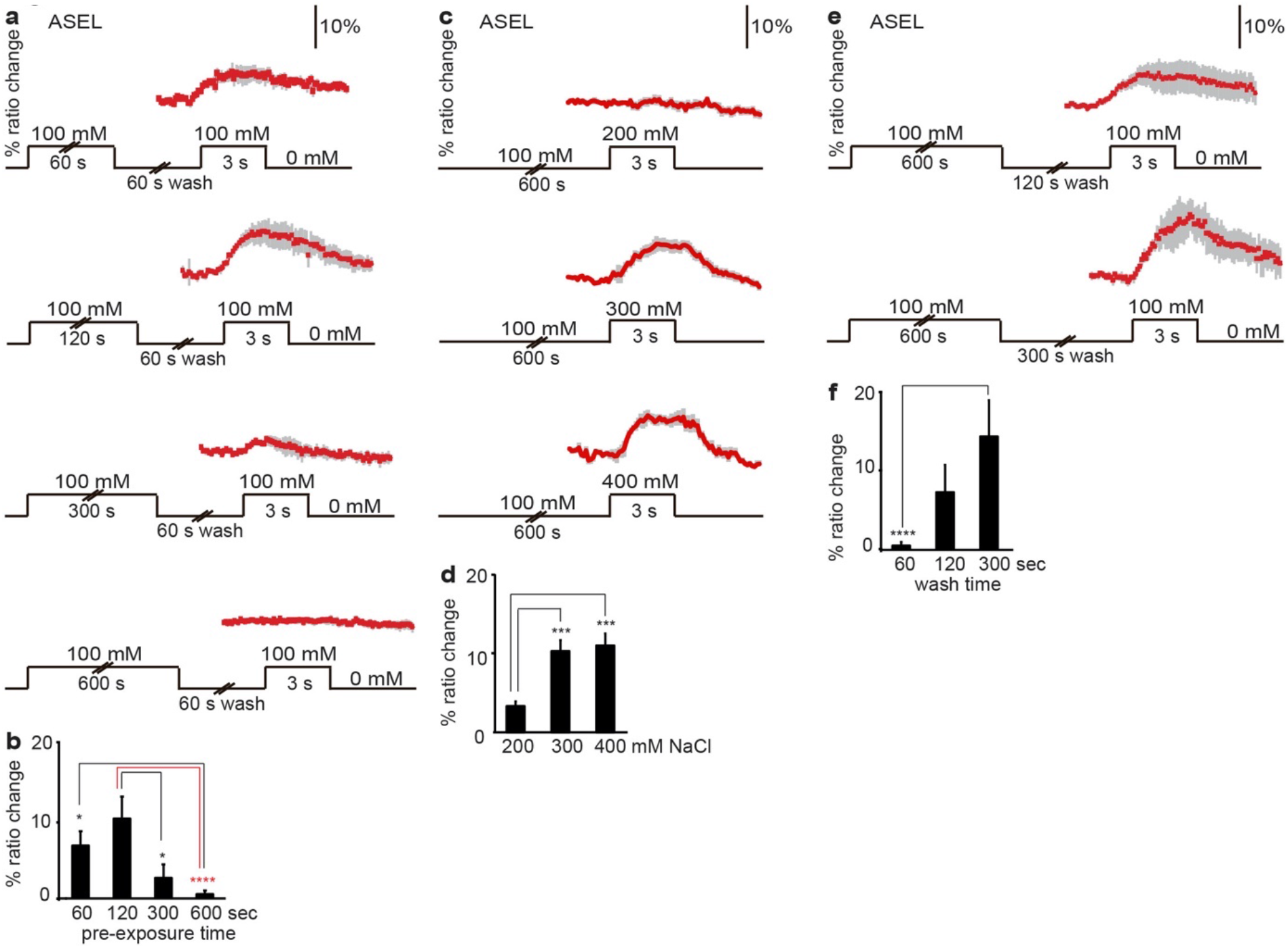
Pre-exposure to NaCl desensitizes ASEL. (**a** and **b**) Animals were pre-exposed to 100 mM NaCl for 60 −600 seconds, washed briefly (60 seconds) and exposed to 100 mM NaCl. (**a**) Average Ca^2+^ transients (± SEM) in ASEL in response to 100 mM NaCl after pre-exposure for 60 seconds n = 13; 120 seconds n = 9; 300 seconds n = 8; 600 seconds n = 16. (**b**) Average maximum ratio changes (± SEM) in ASEL after pre-exposure: responses after 300 or 600 seconds pre-exposure were significantly reduced, compared to responses after 60 or 120 seconds pre-exposure. (**c** and **d**) Animals were pre-exposed to 100 mM NaCl for 10 minutes, before test exposure to 200, 300 or 400 mM NaCl. (**c**) Displays the average Ca^2+^ transients (± SEM) and (**d**) the average maximum ratio changes (± SEM) in ASEL. Exposure to 300 (n=11) or 400 mM (n=7) NaCl yielded significantly stronger Ca^2+^ transients than 200 mM (n = 8). (**e** and **f**) Animals were pre-exposed to 100 mM NaCl for 10 minutes, washed for 1, 2 or 5 minutes in a NaCl-free buffer and re-exposed to 100 mM NaCl, where shows the average Ca^2+^ transients (± SEM) and (**f**) the average maximum ratio changes (± SEM) in ASEL. 2 minutes or longer wash with a NaCl-free buffer restored the Ca^2+^ response of ASEL to 100 mM NaCl. Wash time: 1 minute (n = 16), 2 minutes (n = 8), 5 minutes (n = 6). Statistical significance *: p<0.05, ***: p<0.005, ****: p<0.001 (non-significant differences, p>0.05, have not been indicated).

We further found that ASEL continued to respond to NaCl concentrations above the pre-exposure concentration, e.g. to 300 and 400 mM, but not to 200 mM NaCl (**Fig. 3c,d**). This finding is in accordance with behavioral data that showed that NaCl pre-exposure strongly affected attraction to lower or similar NaCl concentrations but had less or no effect on higher NaCl concentrations (**Supplementary Fig. 2**), suggesting that ASEL desensitization involves threshold modulation.

Adaptation in ASEL is easily reversible, as washing for two or five minutes after 10 minutes of pre-exposure restored reliable Ca^2+^ transients in ASEL (**Fig. 3e,f**; **Supplementary Table 1**). This recovery of the response in ASEL is consistent with behavioral data, where attraction to NaCl after pre-exposure is restored by a 5-minute wash^6^.

We conclude that the ASEL neuron desensitizes with pre-exposure to NaCl and recovers in the absence of NaCl exposure and suggest that this sensory adaptation modulates the strength of attraction to NaCl in behavioral assays.

### Prolonged exposure to NaCl sensitizes ASH

To test if the response of the ASH neurons is affected by pre-exposure, we first exposed animals to 100 mM NaCl for 10 minutes and subsequently introduced a further 100 mM increase to 200 mM NaCl. Strikingly, 16 of the 17 pre-exposed animals (RI 0.94) responded to 200 mM NaCl after pre-exposure, whereas only 5 of 18 animals (RI 0.28) had responded to 200 mM without pre-exposure (**Figure 4a-c**; **Supplementary Table 1**). Response rates and amplitudes to this pre-exposure-stimulus combination were comparable to naïve responses to 500 mM NaCl (**Fig. 4a-c**; **Supplementary Table 1**). Thus, ASH neurons are sensitized by 10 minutes pre-exposure to 100 mM NaCl, upon which they show robust responses to 200 mM NaCl, or to an increase of 100 mM NaCl. Pre-exposure affected neither the number of animals that responded nor the amplitudes of the Ca^2+^ transients in ASH neurons upon exposure to 300 or 500 mM NaCl (**Fig. 4a,c**; **Supplementary Table 1**).

**Figure 4.**
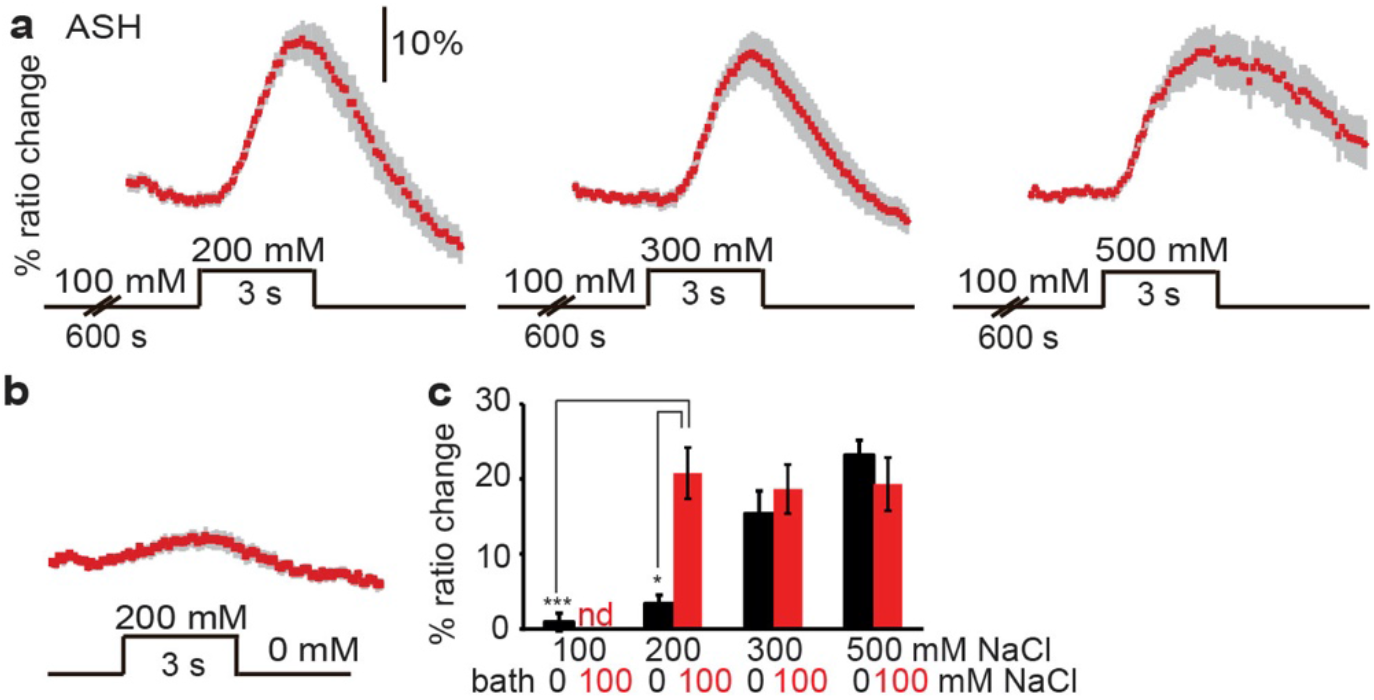
Prolonged exposure to NaCl sensitizes ASH. (**a**) Average Ca^2+^ transients (± SEM) in ASH in response to an increase from 100 mM NaCl (after 600 seconds exposure) to 200 −500 mM NaCl. The response of ASH to 200 mM NaCl was increased after pre-exposure to 100 mM NaCl while the responses to 300 and 500 mM NaCl were unchanged. (**b**) Average Ca^2+^ transients (± SEM) in ASH after exposure to 200 mM NaCl from a baseline of 0 mM NaCl (data from Figure 1). (**c**) Average maximum ratio changes (± SEM) in ASH after exposure to 100, 200, 300, or 500 mM NaCl, in animals pre-exposed to 100 mM NaCl for 600 seconds, or to control condition (100 or 0 mM NaCl bath solution, respectively). Control: 100 mM: n=18, 200 mM: n=18, 300 mM: n=22, 500 mM: n=19. Pre-exposed to 100 mM NaCl: 200 mM: n=17, 300 mM: n=12, 500 mM: n=24. Statistical significance *: p<0.05, *** p<0.005 (non-significant differences, p>0.05, have not been indicated).

### Desensitization of ASEL is likely cell autonomous

Top-down modulation of sensory responses is prevalent in many nervous systems, including in that of *C. elegans*^3^. To determine whether desensitization of ASEL requires input from other neurons, we recorded the Ca^2+^ responses in ASEL neurons of mutants previously shown to affect gustatory plasticity in specific sensory neurons. We tested a mutant in the G protein α subunit *odr-3* that functions in gustatory plasticity in the ADF neurons, serotonergic neurons that play a role in dauer formation and a minor role in chemotaxis to NaCl^2,15,27^. In addition, we tested a mutant in the G protein γ subunit *gpc-1* that functions in gustatory plasticity in the ASI and ASH neurons^6^. ASI neurons are also involved in dauer formation and have a minor role in chemotaxis to NaCl^15^. Furthermore, we tested animals that overexpress the *lsy-6* gene in both ASE neurons, resulting in transformation of the ASER neuron to an ASEL neuron^28^. Finally, we tested *unc-13(e51)* mutants with disrupted synaptic vesicle release, *eat-4(ad819)* mutants with defective vesicular glutamate transport, *unc-31(e928)* and *egl-3(ok979)* mutants with neuropeptide signaling defects, *cat-2(tm2261)* animals with defective dopamine synthesis and *tph-1(mg280)* mutants, which fail to produce serotonin^27,29-33^. Interestingly, none of these mutants showed a significant reduction in desensitization (**Supplementary Fig. 3**), suggesting that desensitization of ASEL is cell autonomous. Only *tph-1* mutants showed slightly abnormal ASEL desensitization; five out of nine animals tested showed a weak response to 100 mM NaCl after 10 minutes pre-exposure, whereas none of the 16 wild type animals tested responded (**Supplementary Fig. 3c**). However, as the average maximum ratio change in pre-exposed *tph-1* animals was not statistically different from that of wild type animals, further analyses are required to reveal a possible contribution of serotonin to ASEL desensitization.

Taken together, our results suggest cell-autonomous desensitization of the ASEL neuron after prolonged exposure to NaCl.

### Sensitization of ASER is likely cell-autonomous

To determine whether sensitization of ASER requires neuropeptide, dopamine or serotonin signaling, we tested the response of the ASER neuron of *egl-3(pk979), cat-2(tm2261)* and *tph-1(mg280)* mutant animals. Mutations in these genes did not affect ASER sensitization (**Supplementary Fig. 4**).

In agreement with previous data^17^, we found a strong response of the ASER neuron of *unc-13(e51)* animals to a decrease in NaCl after 10 minutes exposure, similar to wild type animals (**Fig. 5a,b**), indicating that sensitization of ASER in response to NaCl pre-exposure does not require synaptic neurotransmission. However, unlike the wild type, 67% of *unc-13(e51)* animals also responded to a decrease in NaCl after 30 seconds exposure, resulting in a slow rise of Ca^2+^ (**Fig. 5a-c**). Although these data have not been confirmed in a second *unc-13* mutant strain or by a rescue experiment, our findings suggest that the response of ASER is inhibited by a weak synaptic signal that abolishes the graded depolarization of the cell in naïve (desensitized) animals.

**Figure 5.**
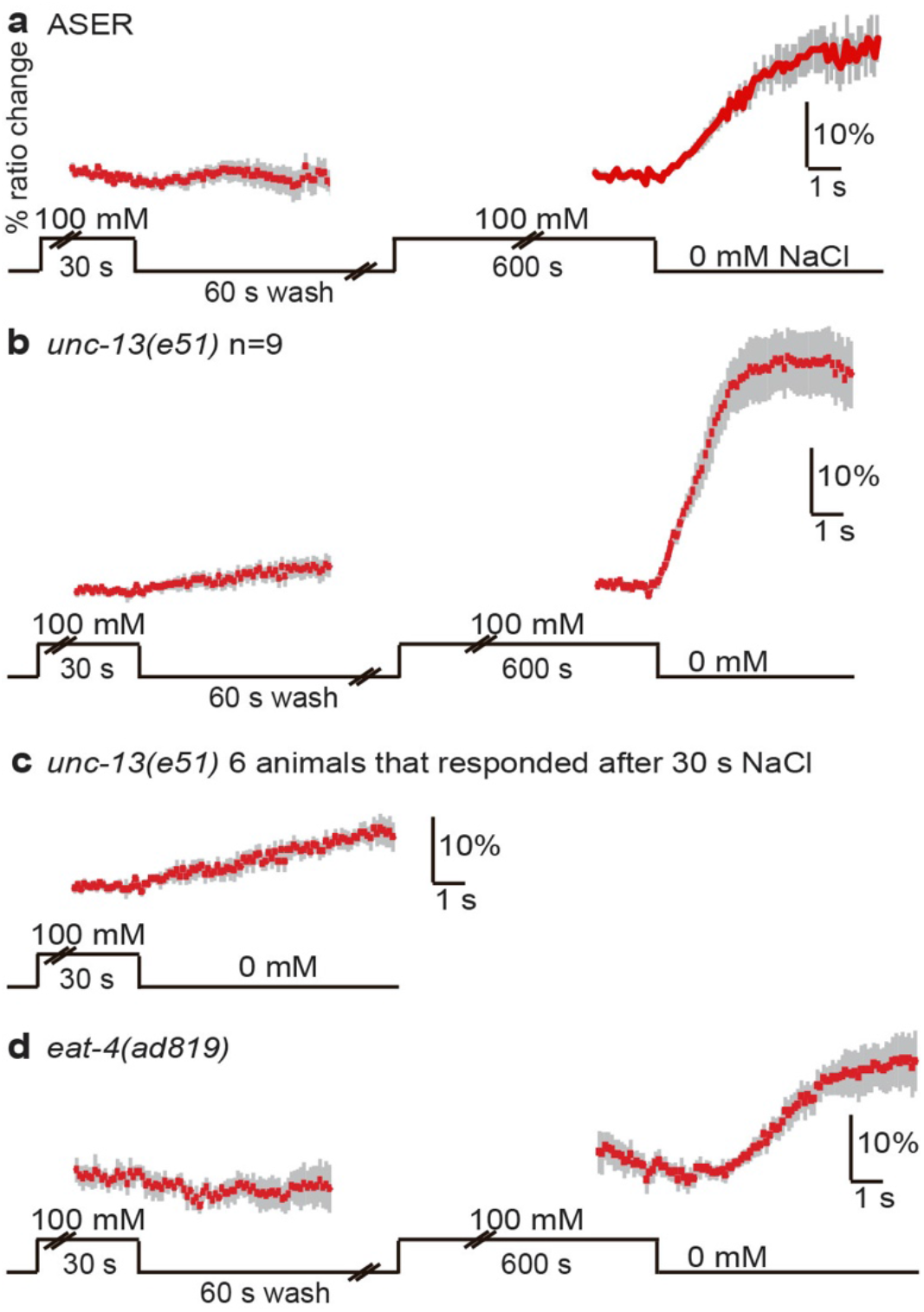
ASER sensitization is affected by *unc-13* and *eat-4* mutations. (**a**) Average Ca^2+^ transients (± SEM) in ASER in wild type animals in response to a decrease in NaCl concentration from 100 mM to 0 mM after 30 and 600 seconds exposure (n=8). (**b**) Average Ca^2+^ transients (± SEM) in ASER of *unc-13(e51)* animals (n=9) in response to a decrease in NaCl concentration from 100 mM to 0 mM after 30 and 600 seconds exposure. 30 seconds exposure to 100 mM NaCl resulted in a small response in ASER, in six out of nine animals tested. Longer exposure to NaCl resulted in a strong Ca^2+^ response of the ASER neurons of *unc-13* animals. (**c**) Average Ca^2+^ transients (± SEM) in ASER of the six *unc-13(e51)* animals that responded to a decrease in NaCl concentration from 100 mM to 0 mM after 30 seconds exposure. (**d**) Average Ca^2+^ transients (± SEM) in ASER of *eat-4(ad819)* animals (n=5) in response to a decrease in NaCl concentration from 100 mM to 0 mM after 30 or 600 seconds exposure. 30 seconds exposure to 100 mM NaCl did not result in a response. 10 minutes exposure to 100 mM NaCl did result in a response, albeit 2 seconds later than in wild type animals.

Finally, *eat-4(ad819)* mutant animals did not respond to a decrease in NaCl after 30 seconds exposure, but did respond after 10 minutes of exposure, indicating that glutamate signaling is not required for ASER sensitization (**Fig. 5a,d**). However, the onset of the ASER response in *eat-4* mutant animals was approximately 2 seconds delayed, compared to the almost immediate response of ASER in wild type animals (**Fig. 5a,d**). Thus, although these findings have not been confirmed by a rescue experiment, glutamate seems to be involved in facilitating a rapid onset of a Ca^2+^ response to a decrease in NaCl. Since no delay in ASER response was observed in *unc-13(e51)* mutant animals, the glutamate signal might be extra-synaptic in origin. Further experiments are required to reveal the nature of this signal.

Taken together, we conclude that sensitization of the ASER neuron in response to prolonged exposure to NaCl is likely cell-autonomous. We also found that the ASER Ca^2+^ response likely involves a glutamate-mediated signal that facilitates a rapid onset of the response, and a synaptic signal that abolishes a weak graded response in naïve animals, neither of which are required for gustatory plasticity.

### ASH sensitization requires ASE, glutamate, neuropeptides, dopamine and serotonin

Since ASE neurons are required for gustatory plasticity^2^ we tested whether they are required for sensitization of ASH by measuring ASH responses in *che-1* mutants that lack functional ASE neurons^34^. *che-1* mutants were indistinguishable from wild type animals in their naïve response to 200 or 500 mM NaCl (**Fig. 6c,e**, compare to wild type naïve responses in **Fig. 1d,e**), but failed to respond or responded very weakly to 200 mM NaCl following pre-exposure to 100 mM NaCl for 10 minutes (**Fig. 6a,d,e**). To control for the responsiveness of the ASH neurons of the tested animals, we confirmed the response of the same animals to 500 mM NaCl (**Fig. 6d**). These data show that ASH sensitization is a circuit effect that requires the ASE neurons. Whether ASE neurons recruit ASH by allowing it to sense NaCl at lower concentrations (akin to a form of threshold modulation), or whether an ASEL/R sensory signal is transmitted to ASH that effectively acts as an interneuron, reminiscent of AWC recruitment by ASE^35^, remains unknown.

**Figure 6.**
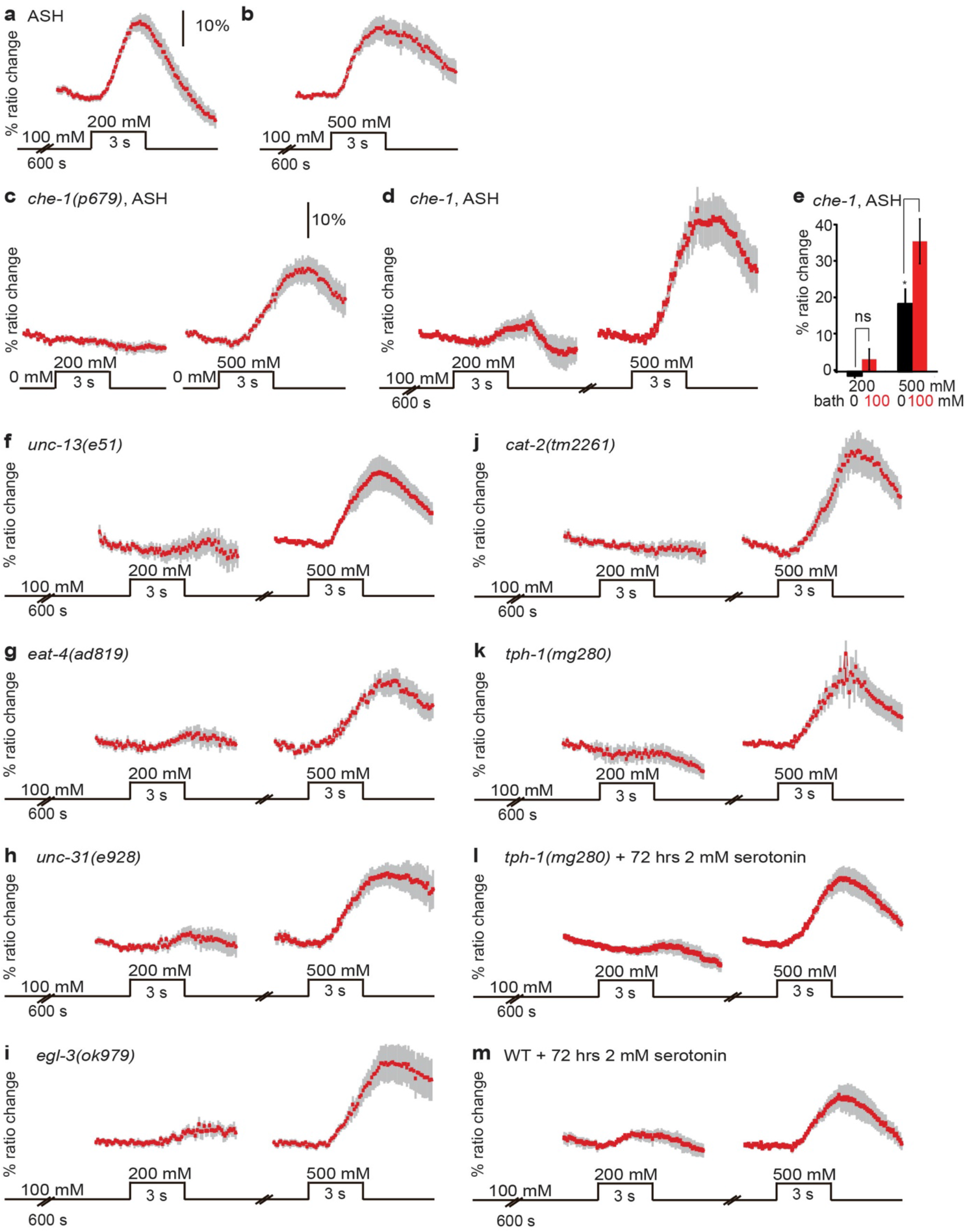
ASH sensitization requires ASE, glutamate, neuropeptides, dopamine and serotonin. (**a**,**b**) Average Ca^2+^ transients (± SEM) in ASH neurons of wild type animals after 600 seconds pre-exposure to 100 mM NaCl in response to an increase to 200 (**a**) or 500 (**b**) mM NaCl (data from Figure 4). (**c**) Average Ca^2+^ transients (± SEM) in ASH neurons of *che-1(p679)* animals in response to exposure to 200 or 500 mM NaCl. Naïve *che-1* animals did not respond to 200 mM NaCl (n = 6) but did respond to 500 mM (n = 11). (**d**) Average Ca^2+^ transients (± SEM) in ASH neurons of *che-1(p679)* animals after 600 seconds pre-exposure to 100 mM NaCl in response to an increase to 200 or 500 mM NaCl. Only very weak responses to 200 mM NaCl could be observed in *che-1* animals (n = 11, 3 out of 11 worms showed a response). The same *che-1* animals did respond to 500 mM NaCl after a 2 minutes wash (n = 11). (**e**) Average maximum ratio changes (± SEM) in ASH of *che-1* mutant animals after exposure to 200 or 500 mM NaCl, in animals pre-exposed to 100 mM NaCl for 600 seconds, or to control condition (100 or 0 mM NaCl bath solution, respectively). (**f-m**) Average Ca^2+^ transients (± SEM) in ASH neurons of various neurotransmitter mutants after 600 seconds pre-exposure to 100 mM NaCl in response to an increase to 200 and subsequently 500 mM NaCl. None of these mutants showed a significantly stronger response to 200 mM NaCl after pre-exposure than in the naïve situation (p>0.05). Only responses of animals that responded to 500 mM NaCl were included. (**f**) Two out of eight *unc-13(e51)* animals tested responded to 200 mM NaCl after pre-exposure to 100 mM NaCl. (**g**) Two out of nine *eat-4(ad819)* animals responded. (**h**) Three out of seven *unc-31(e928)* animals responded. (**i**) Three out of nine *egl-3(ok979)* animals responded. (**j**) Three out of nine *cat-2(tm2261)* animals responded. (**k**) One out of eight *tph-1(mg280)* animals showed sensitization of ASH. (**l**) Four out of ten *tph-1* animals cultured in the presence of 2 mM serotonin responded to 200 mM after exposure to 100 mM NaCl. (**m**) Four out of six wild type animals cultured in the presence of 2 mM serotonin responded to 200 mM after exposure to 100 mM NaCl. Statistical significance ns: p>0.05, *: p<0.05, *** p<0.005 (non-significant differences, p>0.05, have not been indicated).

We next asked which neurotransmitters play a role in ASH sensitization. Interestingly, all mutants tested, *unc-13(e51), eat-4(ad819), unc-31(e928), egl-3(ok979), cat-2(tm2261)* and *tph-1(mg280)*, showed reduced sensitization of ASH, but responded as wild type to 500 mM NaCl, suggesting that synaptic transmission, glutamate, neuropeptides, dopamine and serotonin signaling all play a role in sensitization of ASH (**Fig. 6f-k**). Previously, Hilliard *et al*. have shown that mutation of *unc-13* or incubation in serotonin does not affect the Ca^2+^ response of ASH to an osmotic stimulus^23^.

To confirm the contribution of serotonin to sensitization of ASH, we cultured *tph-1* mutant animals for 72 hours on plates containing serotonin. Supplying either 2 or 4 mM serotonin did not significantly restore ASH sensitization in these animals (**Fig. 6l**), but reduced sensitization of wild type animals (**Fig. 6m**), suggesting that serotonin levels need to be precisely regulated to sensitize ASH. Serotonin may have a further role during development^5^. We conclude that the recruitment of ASH to respond to non-toxic levels of NaCl requires ASE and relies on multiple pathways involving synaptic transmission, glutamate, neuropeptides, dopamine and serotonin signals.

### *In silico* sensory neurons support fast dynamic range adaptation to NaCl

To better understand the behavioral implications of the different forms of sensitization and desensitization in ASEL, ASER and ASH neurons, we used our empirical results to construct a computational model (**Fig. 7a-d**). Many *C. elegans* sensory neurons, including ASE and ASH, respond to the change in stimulus over time^17,36^. Transient pulse-like responses are well captured by two opposing and timescale separated components^37^. In the absence of detailed conductances, we imposed dynamics that closely mimic models of eukaryotic chemotaxis^38^ (**Fig. 7a-c**; Supplementary methods) by letting the slow variable, hyperpolarizing current, denoted *S*, follow the fast, depolarizing current, *F*, with a delay (**Fig. M2** in Supplementary methods). The model ASEL depolarizes to NaCl increases, while ASER depolarizes to NaCl decreases and hyperpolarizes to NaCl increases^17^. Since ASHL and ASHR respond identically to NaCl and are electrically coupled, we modeled them as a single unit which depolarizes to NaCl increases (ASH, **Fig. 7c**)^36,39-41^. As hyperosmotic responses are unaffected by gustatory plasticity, they were not considered in our model.

**Figure 7.**
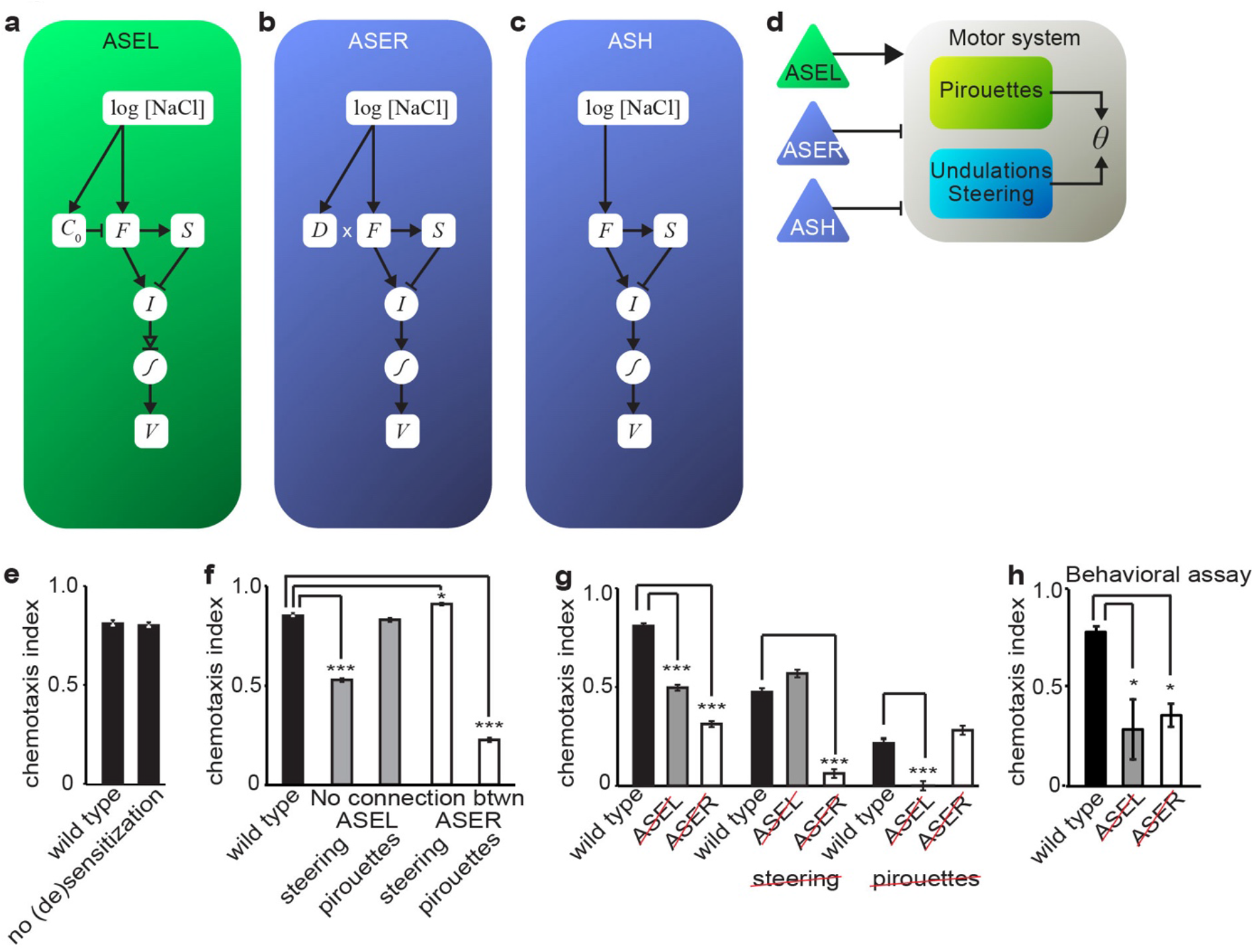
Computational model reproduces sensory responses to NaCl in virtual quadrant assay. (**a-d**) Schematic of the computational model. (**a-c**) Internal sensory computation in ASEL, ASER and ASH, respectively. Sensory stimuli drive a fast component *F*, and a delayed rectifier *S*, with opposite contributions to the overall current *I*. ASEL desensitization is included as an adaptive threshold *C*_*0*_, ASER sensitization is modeled using a multiplicative gain, *D*, and ASH sensitization is modeled as a stochastic binary switch. The current-voltage relation is given by a sigmoidal activation function. (**d**) Sensorimotor pathway modulates rhythmic undulations and the frequency of random turns. (**e-g**) Average chemotaxis index (± SEM) of model WT versus mutant animals in the simulated quadrant assay. (**e**) Our model shows no difference in attraction in the quadrant assay between wild type model animals and animals with always fully sensitized ASEL and ASER (no (de)sensitization). Ablating the synaptic connections from ASEL to the steering circuit or from ASER to the pirouette neuron significantly reduced the chemotaxis index (**Table M1**). Ablating the connection from ASER to steering very slightly increased the chemotaxis index. (**g**) “Double mutants” with ablated ASEL (ASEL) and no steering (steering), or ablated ASER (ASER) and no pirouettes (pirouettes) achieved the same chemotaxis index as single mutants. (**h**) Experimental validation: animals with genetically ablated ASEL (OH8585 ASEL) and genetically ablated ASER (OH8593 ASER) exhibited a strong reduction in the average chemotaxis index (± SEM) in the quadrant assay. See corresponding **Supplementary Movies S1-S14**. *: p<0.05, ***: p<0.0005 (non-significant differences, p>0.05, have not been indicated).

**Figure 8.**
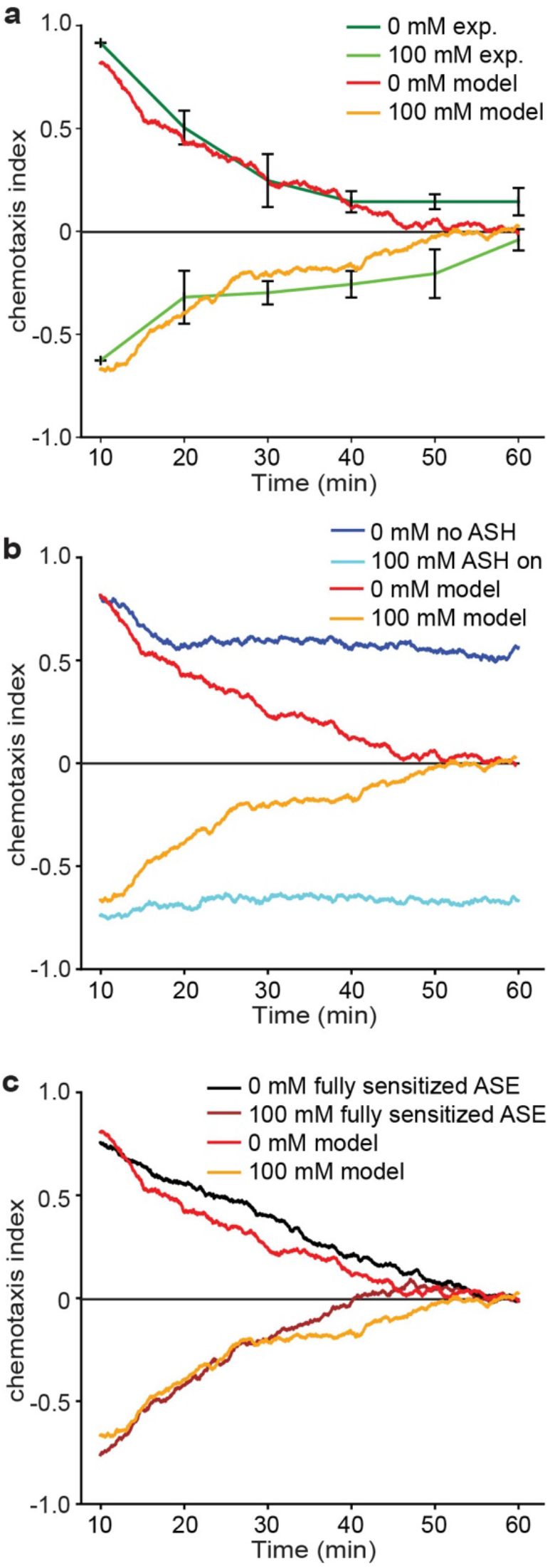
Stochastic recruitment of ASH drives gustatory plasticity in our computational model. Chemotaxis index over time in experiments and simulations of the quadrant assay. (**a**) Average chemotaxis index (± SEM) of naïve (0 mM) and pre-exposed (15 minutes, 100 mM NaCl) animals. The behavioral results (green) and the modeling results (red, orange) show a monotonic decay towards a chemotaxis index of 0, for both naïve and pre-exposed animals. (**b**) Virtual chemotaxis index for animals with unrecruited ASH (dark blue) and with recruited ASH (light blue). Without ASH state dynamics the chemotaxis index does not decay to 0. (**c**) Comparison of chemotaxis index for model with ASE (de)sensitization (red, orange) and with fully sensitized ASEL and ASER (black, brown).

We parameterized the model sensory cells using Ca^2+^ imaging data from fully sensitized ASEL and ASER neurons. The rise times, and even more so, the decay times of the responses were consistently and significantly faster in ASEL than in ASER, consistent with previous work^17^ (**Fig. 1a** and **2a**; **Supplementary Table 1**; **Supplementary Fig. 5**; decay times: ASEL responses were back to baseline in 3.9 ± 1.3 seconds, whereas ASER responses decreased by approximately 15% of the maximum amplitude in 6 seconds).

Our and previous^17^ Ca^2+^ imaging results consistently show that the peak depolarization amplitude varies with the stimulus intensity. The attractive chemotaxis responses to NaCl concentrations ranging from 0.1 mM to 100 mM best fit a logarithmic relationship^2,6^. Similarly, we found that a logarithmic function of stimulus intensity best reproduced our ASEL and ASER Ca^2+^ response data, strongly reminiscent of the Weber-Fechner law of sensory perception, which states that the ability to distinguish between two magnitudes of a stimulus scales with the magnitude; a mathematically equivalent form is, *R* ∝ log *S*, where *R* and *S* are the response and stimulus, respectively^3,19^. We chose a parsimonious representation of this dynamic range modulation in which sensory neurons instantaneously respond to the logarithm of the NaCl concentration (Supplementary Methods, **Fig. 7a-c**). Results showed close agreement with Ca^2+^ traces in ASEL and ASER (**Fig. M3** in Supplementary methods).

### Gustatory adaptation occurs downstream of dynamic range adaptation

We next incorporated adaptation into our model sensory neurons, with parameters constrained by our Ca^2+^ imaging data. ASEL desensitized to stimuli below the concentration of NaCl pre-exposure, but continued to respond to higher concentrations, consistent with threshold adaptation (*C*_*0*_ in **Fig. 7a**). ASER adaptation was modeled as gain modulation (*D* in **Fig. 7b**), consistent with the absence of a response in the naïve context (**Fig. 1c**). Both ASEL and ASER (de)sensitization had to be applied after logarithmic scaling to reproduce the Ca^2+^ imaging data. Thus, this model constraint suggests that gustatory adaptation occurs downstream of the receptor and of rapid dynamic range adaptation.

Our Ca^2+^ imaging data indicate that ASH is recruited into the low-concentration NaCl sensing circuit upon pre-exposure to NaCl. In the absence of mechanism, we modeled this minimalistically as an on/off switch, recruiting and releasing ASH from the gustatory circuit. Switching is governed by a stochastic process with dynamic switching rates dependent on the history of the salt concentration (see Supplementary methods).

### *In silico* animals reproduce neuronal response and behavior of naïve animals

To model the behavioral consequences of sensory adaptation, we constructed a full sensory-motor model that could be simulated in a virtual assay arena. As we focused on the sensory system, we chose to use a minimal embodiment and a relatively abstract motor system (**Fig. 7d**). To explore and navigate their environment, *C. elegans* use a combination of steering, where animals gently turn to reorient, and a biased random walk, in which animals reorient by making sharp turns or pirouettes^16,26^. Both strategies allow animals to migrate along chemical gradients. In our model, point animals moved at a fixed speed of 0.11 mm/sec^36^ with a dynamic bearing, subject to both steering and pirouettes (see Supplementary methods).

To explore gustatory plasticity *in silico*, we replicated the quadrant NaCl-choice assay^6,42^ in our model. Simulations of 1000 wild type naïve worms, with ASEL, ASER and ASH adaptation/recruitment dynamics, for ten minutes of virtual time in the quadrant assay, yielded a similar *in silico* chemotaxis index to experimental naïve results (wild type in **Fig. 7e,h**).

### ASE (de)sensitization reduces robustness of chemotaxis in our computational model

The opposite actions of ASEL and ASER adaptation suggest only one of the ASE pair is fully sensitized at any one time. Thus, we expected a severe performance penalty in our simulations, relative to a model with no adaptation. We found that both models with and without sensory adaptation in ASE quantitatively reproduce the chemotaxis index from the quadrant assay (**Fig. 7e; Supplementary Movies S1 and 2**). To better understand the ramifications of adaptation on performance and robustness, we generated population models of animals, which we then simulated on our choice assay. Each population consisted of model animals either with (test) or without (control) ASE adaptation. To focus on the role of adaptation, within each population, model animals differed only in their ASE kinetic parameters (determining the rise and decay profiles; see Supplementary Methods). Across a wide range of noise amplitudes, and hence a wide range of sensory neuron parameters, model worms with ASE (de)sensitization performed at least as well as those without sensory adaptation (**Supplementary Fig. 7a**). Thus, our computational model predicts that the robustness of the performance in the quadrant assay (as measured by the chemotaxis index) is not reduced by ASE (de)sensitization.

### ASH Sensitization reproduces *in silico* gustatory plasticity

Next, we looked at the behavior of pre-exposed animals focusing on a 15 minutes 100 mM NaCl pre-exposure, at which the avoidance behavior is the strongest (**Supplementary Fig. 2**). Since ASER sensitizes, producing stronger attraction, and ASEL desensitizes, producing weaker attraction but not avoidance, ASH seemed a likely candidate to drive salt avoidance. Indeed, to reproduce strong avoidance after pre-exposure, we had to set the synaptic weights of ASH to be significantly stronger than the ASE synaptic weights (**Fig. 9a**, 10 min., **Table M1** in Supplementary methods). Such a ‘drowning’ of attractive signals by ASH is consistent with the strong Ca^2+^ response in ASEL to 300 and 500 mM NaCl in wild type animals (**Fig. 1a**) and with behavioral results: While wild type *C. elegans* are strongly repelled by these concentrations, ASH deficient *odr-3(n1605)* animals are strongly attracted by them^2^. In addition, genetic ablation of ASH strongly reduced avoidance after pre-exposure^2^, consistent with the results of our computational model.

**Figure 9.**
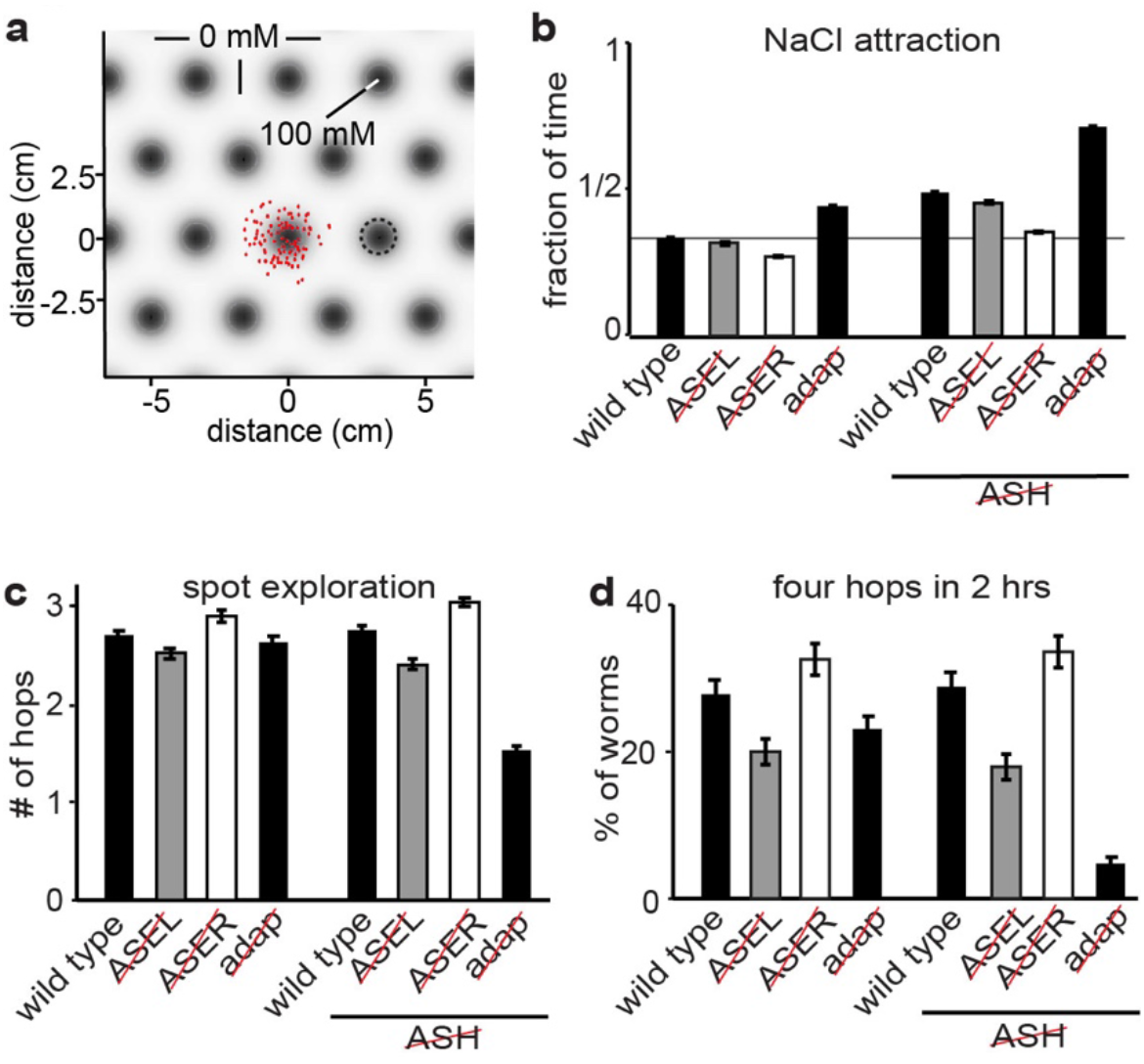
Sensory adaptation controls exploration in a virtual spot assay. (**a**) Spot assay arena with identical salt spots with standard normal salt concentration peaking at 100 mM. Peaks are positioned on an infinite hexagonal grid. The standard deviation for each spot is 0.6 cm and the distance between two nearest spots is 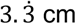. Populations of 500 worms each were simulated for 1 hour. Naive worms were initialized in the center of a single spot, with random orientations. (**b**) Salt attraction in the spot assay for eight different populations, measured as the fraction of time a worm dwells on salt (>25 mM), averaged over the population (± SEM). (**c, d**) Exploration quantified as the number of times a worm changes from one spot to another for eight different populations. For each population, 500 worm trajectories were analyzed. (**c**) The average number of hops achieved within 2 hours (± SEM). (**d**) The percentage of worms that made four or more hops within 2 hours (± SEM). Note the 6-fold difference between the simulations with (‘wild type’, 28.8%) and without (‘adap’, 4.8%) in populations of model animals lacking ASH.

### Sensory neuron timing strongly influences navigation strategies in our model

Our and published Ca^2+^ imaging experiments^17^ have revealed a significant timescale separation between the Ca^2+^ responses of fully sensitized ASEL and ASER, both in their rise and decay times (**Supplementary Fig. 5**). To determine the behavioral consequences of the timescales of ASEL and ASER kinetics, we determined the contributions of ASEL and ASER to steering and pirouettes in our simulations. When we ablated the *in-silico* connection from ASEL to the pirouette motor program or the *in-silico* connection from ASER to the steering circuit, chemotaxis remained unchanged relative to wild type model animals (**Fig. 7f; Supplementary Movies S1, S3-S6**). Conversely, virtually severing the connection from ASEL to the steering circuit or from ASER to the pirouette motor program severely reduced chemotaxis (**Fig. 7f; Supplementary Movies S1, S3-S6**). Thus, in our model, ASEL controls steering, but has little effect on the pirouette rate, whereas ASER modulates pirouettes, but has little control over steering. We found that disabling steering in simulations of wild type and ASEL-ablated animals equally reduced chemotaxis in the virtual quadrant assay (**Fig. 7f; Supplementary Movies S1, S7-S14**), confirming this model behavior. Similarly, disabling pirouette modulation in wild type or ASER ablated animals resulted in equally reduced chemotaxis (**Fig. 7f; Supplementary Movies S1, S7-S14**). Finally, ablating ASEL in animals where pirouette modulation was disabled, or ablating ASER in animals where steering was disabled almost fully abolished the response to NaCl in our model (**Fig. 7f; Supplementary Movies S1, S7-S14**). These results are in agreement with behavioral results of Suzuki et al. who previously showed that ASEL activation promotes runs whereas ASER activation induces turns^17^.

Our computational model predicts that both ASEL and ASER contribute to chemotaxis in the quadrant assay. To test this, we genetically ablated either ASEL or ASER, using animals that express Caspase-3 in either the left or right ASE neuron^24^. These animals showed strongly reduced chemotaxis to NaCl (**Fig. 7h**), confirming that both ASEL and ASER contribute to navigation in the quadrant assay.

To rule out any contribution of desensitization of ASEL and sensitization of ASER to the above analyses, we re-ran our simulations with both ASE neurons fully sensitized and ASH recruitment disabled. These analyses gave results very similar to our previous analyses (**Fig. 7g**; **Supplementary Fig. 7b**), confirming that in our model the separate roles of ASEL and ASER in motor control are direct consequences of the rise and decay times of their responses.

Thus, our model results are consistent with results from ASEL and ASER ablated animals, as well as with observations that both steering and pirouettes contribute to the chemotaxis index in the quadrant assay. In the model, distinct motor programs are separately controlled by ASEL and ASER, as a direct result of the timescales of the sensory neuron responses. To steer, sensory signals must be detected on the timescale of a half-undulation: *O*(1-2 sec) or faster^26,43,44^ (**Fig. M4** in Supplementary methods). The slower rise time in ASER precludes this (**Supplementary Fig. 7b**). The contribution of ASER to steering in different assays^26,45^ may indicate a faster rise time in ASER. Pirouettes occur with a mean rate of 2.1 events per minute^26^ or less (in the quadrant assay). Therefore, to effectively modulate this rate requires a memory of salt exposure over commensurate (or longer) timescales. The fast decay time of ASEL precludes this, while the slow decay time of ASER is ideally suited to modulate the pirouette rate effectively. Should ASER decay on a faster timescale, the modulation of pirouettes would require a slow integration elsewhere in the circuit.

### *In silico* ASH mediates a detailed balance between attraction and avoidance

We next asked whether sensory adaptation is sufficient to account for the balance of attraction and avoidance over time. We therefore followed the long-term behavior of real animals in the quadrant assay over one hour. For both naïve and pre-exposed animals, the chemotaxis index dropped to approximately zero over the course of the hour, indicating roughly equal numbers of animals in the salt and no salt quadrants (**Fig. 8a**, green lines). Strikingly, without any further parameter tuning, simulations of naïve and pre-exposed animals closely reproduced these experimental data (**Fig. 8a**, red and orange lines).

To determine the potential contribution of ASH sensitization to the chemotaxis index decay to zero over time we simulated animals having either ASH completely disabled or fully recruited. Now the chemotaxis index decayed only partially, reaching a plateau around 0.6 and −0.7 respectively (**Fig. 8b**, blue lines). Conversely, our simulations including ASH sensitization dynamics showed that ASH recruitment inside the NaCl quadrants (driving avoidance) and ASH relaxation outside the NaCl quadrants (allowing attraction) lead to an equal number of animals inside and outside the NaCl quadrants. These results point at a detailed balance description^46^ of ASH dynamics, in which animals stochastically switch between recruited and unrecruited ASH states, consistent with the finding that on average ASH dynamics are governed by similar time scales of sensitization and de-sensitization. Our simulations of naïve animals with and without ASE adaptation yielded similar chemotaxis indices in the quadrant assay over 10 minutes (**Fig. 7e**) and over an hour (**Fig. 8c**, black and brown lines).

In summary, our model predicts that dynamic-state switching of ASH mediates the behavioral switch associated with gustatory plasticity. Neither desensitization of ASEL nor sensitization of ASER appear to play a role in the quadrant assay, aside from their possible contribution to the recruitment of ASH.

### Sensory adaptation enhances exploration in an *in silico* salt spot assay

The absence of an obvious role for ASE sensitization in the virtual quadrant assay raises questions about the possible benefit of ASE adaptation in *C. elegans*. We conjectured that ASEL and ASER adaptation may serve to modulate attractive search behaviors in more natural, heterogeneous environments. To test this, we constructed a virtual spot assay consisting of an infinite grid of identical spots with radial Gaussian concentration profiles (**Fig. 9a**). Model animals were placed in the center of one spot from which they were free to move up and down salt gradients, allowing them to visit different salt spots. Attraction to NaCl of the model animals was quantified as the fraction of time spent on NaCl spots (>25 mM), averaged over the population (**Fig. 9b**). Exploration behavior was quantified in terms of hops from one spot to another (**Fig. 9c,d**).

In this spot assay, model animals displayed a balance between localized attraction to NaCl and exploratory behavior (**Supplementary Movie S15**). Without ASH recruitment, adaptation-defective animals in which ASEL and ASER were fully sensitized (“adap” in **Fig. 9**) exhibited stronger NaCl attraction, that led to the majority of animals remaining very close to their initial location for the duration of the simulation (**Fig. 9b-d, Supplementary Fig. 8, Supplementary Movie S15-30**). Incorporating ASEL and ASER (de)sensitization resulted in reduced local NaCl attraction and enhanced exploratory behavior (**Fig. 9b-d, Supplementary Fig. 8**). Enabling ASH recruitment further enhanced exploratory behavior (**Fig. 9b-d, Supplementary Fig. 8**). In our model, ASEL desensitization enhanced exploration during salt attractive behaviors by increasing the typical exploration radius of a spot (and hence the rate of escape from a given spot), ASER sensitization limited the attractive response to sufficiently large spots (or sufficiently long dwell-times on a spot), whereas ASH recruitment led to more widespread dispersal of the population (**Fig. 9d, Supplementary Fig. 8**).

Next, we determined the relative contributions of ASEL and ASER to exploration. We found that removing ASEL reduced exploration, whereas removing ASER in our virtual animals increased exploration (**Fig. 9c**,**d, Supplementary Fig. 8**). Unlike the quadrant assay, however, the contribution of ASEL to NaCl attraction was limited in the spot assay, whereas removing ASER had a stronger effect (**Fig. 9b**).

## Discussion

In naïve *C. elegans*, attractive, ASE mediated, and aversive, ASH mediated, salt responses are controlled by clearly delineated subcircuits, resulting in a switch between attraction up to 200 mM NaCl and avoidance of higher concentrations (Fig. 10). However, while naïve animals are attracted to low salt concentrations, extended NaCl exposure without food leads animals to avoid any NaCl concentration^2,5,6^, implying an adaptive foraging behavior. Here, we showed that the behavioral switch between NaCl attraction and avoidance is mediated by plasticity in sensory neurons, resulting in altered dynamic ranges in both attractive and nociceptive subcircuits. Based on our experimental and modeling results we propose that the sensory response of *C. elegans* to NaCl is regulated at multiple levels.

**Figure 10.**
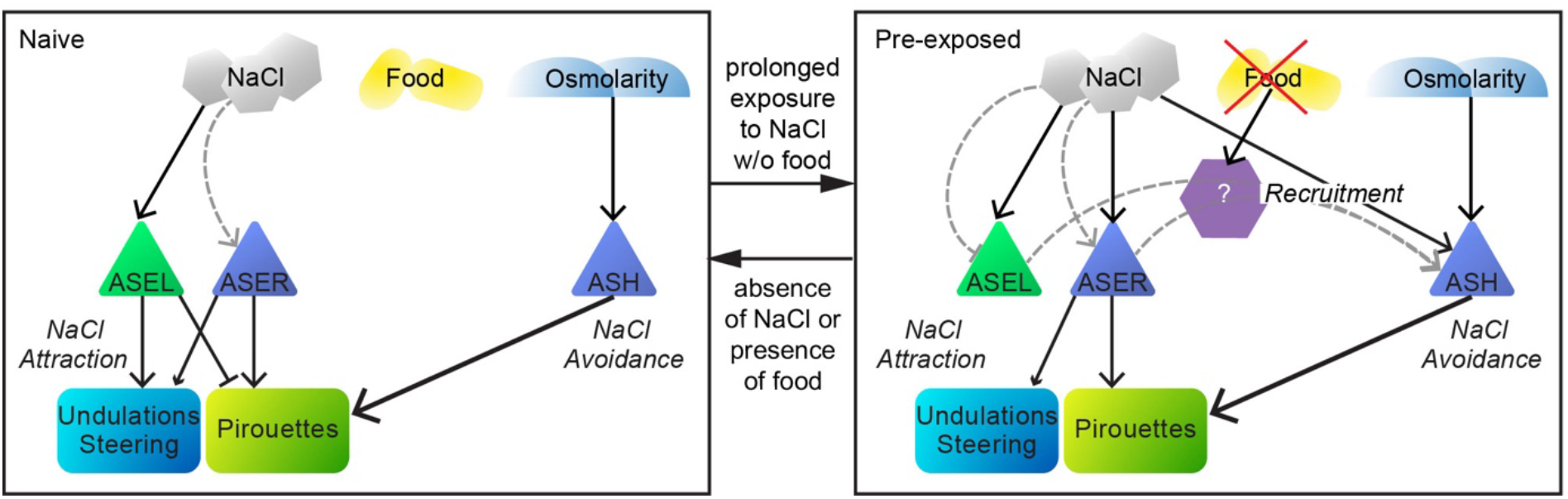
Schematic model of the NaCl navigation circuit. Schematic of the different forms of sensory adaptation and their downstream effects in response to NaCl exposure in the absence of food. Left: The naïve state, in the absence of NaCl and/or the presence of food. ASEL is fully sensitized, ASER desensitized and ASH only responds to high NaCl concentrations (osmotic shock). Right: pre-exposed state, after 10-15 minutes of exposure to NaCl in the absence of food: ASEL becomes desensitized, ASER sensitized and ASH recruited to respond to lower NaCl concentrations. Recruitment of ASH depends on an absence of food signal and ASE, possibly via one or more intermediate neurons. It is unclear whether an absence of food signals recruits ASH (as indicated schematically in the figure), or whether the presence of food inhibits the recruitment of ASH. ASEL and ASER mediate attraction to NaCl and ASH mediates avoidance of NaCl. NaCl dependent adaptation presented in gray dashed arrows. Solid arrows represent excitation (either via receptors or synapses), solid bars inhibition.

We observed an effective flip between the dynamic ranges of the primary avoidance neurons, ASH, and salt attractive neuron, ASEL, which suppresses attraction and enhances avoidance of naïvely attractive NaCl concentrations. Surprisingly, ASER, the second primary sensor mediating NaCl attraction sensitizes with NaCl-exposure. Hints of opposite forms of adaptation in ASER and ASEL were already reported by Oda *et al*., who found adaptation after 10 minutes of exposure to 20 mM NaCl^25^. Our analyses suggest that both ASEL and ASER (de)sensitization are mostly cell autonomous. In contrast, sensitization of ASH requires signals from the ASE neurons, glutamatergic, serotonergic, dopaminergic and neuropeptide signaling, underscoring the complexity of this seemingly simple behavioral paradigm. One of these signals probably mediates the cue that food is lacking. Candidate cells for detecting the absence of food are the ASG pair of amphid sensory neurons, as these have been shown to play a similar role in taste avoidance learning^47^. However, it remains unclear whether absence of food signals recruit ASH and/or whether the presence of food inhibits the recruitment of ASH. Accordingly, our model remains agnostic to these possibilities, as the encoding of food in the model is implicit.

While multiple sensory neurons and contributions from the downstream circuitry most likely contribute to the rich behavioral responses in *C. elegans*, our simulations demonstrate the feasibility of a parsimonious model in which recruitment of ASH by an ASE (NaCl) derived signal underpins the switch between attraction and avoidance in gustatory plasticity.

The bilaterally asymmetric kinetics of the ASE neurons, in which ASEL and ASER depolarize in response to concentration increases and decreases, respectively^17^, suggest a potential to double the ASE sensory dynamic range (from {*0<,x*} to {*-x<,x*}), thus enhancing the resolution of NaCl sensing in the animal. However, the apparent timescale separation in the responses suggests otherwise. Our computational model demonstrates how a separation of timescales leads to distinct pathways that control different motor programs. ASEL and ASER have previously been linked to the control of steering and pirouettes, respectively^17^. In our model, fast sensory processing in ASEL controls steering, whereas slow sensory processing in ASER modulates pirouettes over tens of seconds or minutes. Such encoding of distinct motor actions in the kinetics of neuronal activation provides an effective mechanism for dictating behavioral output at any point along the sensory-motor pathway, even in the sensors themselves.

While one would expect some forms of adaptation in sensory neurons^3,48^, our results point to severe information loss, causing a potentially considerable impediment in salt sensing. When ASER is desensitized the animal’s ability to respond to concentration decreases is almost abolished. Our model predicts that naive worms (with sensitized ASEL only) will move up gradients. If the NaCl regions traversed are insufficient to sensitize ASER, desensitization of ASEL will promote dispersion from NaCl-rich regions. Conversely, if ASER is sensitized, a trajectory down the NaCl gradient will be suppressed by promoting turning. Combined, ASE (de)sensitization would ensure that animals only respond to sufficiently large NaCl regions, ignoring small fluctuations. In summary, NaCl-adaptation of ASE neurons could serve to balance exploration and exploitation navigational strategies in complex, heterogeneous environments, as in our virtual spot assay.

In addition, if ASEL predominantly controls steering towards gradient peaks when navigating up the gradient, desensitization would reduce steering only after entering a salt region (promoting broader exploration within the region). Conversely, if ASER predominantly modulates pirouettes, then leaving a salt patch will likely induce a pirouette. The above reasoning is consistent with our model assumption that ASER mediates attraction only, an effect that is masked when ASH is recruited. In addition, ASER could mediate avoidance in gustatory plasticity by flipping its synaptic sign to a downstream interneuron in a food/starvation dependent way^49^.

*C. elegans* feed on bacteria in patchy environments, most densely in rotting vegetation. These bacterial patches likely vary in size and may be well separated spatially, consistent with the *C. elegans* “boom-and-bust” life cycle^50^. As food is depleted, animals disperse and forage, often through highly variable and uncertain environments, which call for random foraging strategies. Close enough to a signal, active steering and turning are beneficial. Strategies involving balance of exploration and exploitation make sense in this context.

The intuition presented here suggests that the compact nervous system of *C. elegans* may benefit from enhanced computation in sensory neurons at the price of considerable information loss. Taken together, our computational model and our and previous experimental data point to a highly complex set of distinct forms of plastic sensory computation in the NaCl sensing circuit, indicating that, compared to higher animals, *C. elegans* has seen a shift of computation from the inter-to sensory layers over its evolutionary history.

## Methods

### Strains and germline transformation

The following strains were used in this study

Wild-type *C. elegans* strain used was Bristol N2.

GJ243 *che-1(p679)I; gjEx513[sra-6::YC3*.*60 elt-2::GFP]*, 0x outcrossed

GJ254 *gjEx523[sra-6::YC3*.*60 elt-2::GFP]*

GJ282 *gpc-1(pk298)X; gjEx549[sra-6::YC3*.*60 elt-2::GFP]*, 6x outcrossed

GJ285 *gpc-1(pk298)X; gjEx552[flp-6::YC3*.*60 elt-2::GFP]*, 6x outcrossed

GJ1494 *odr-3(n1605)V; gjEx866[flp-6::YC3*.*60 elt-2::GFP]*, 6x outcrossed

GJ1497 *tph-1(mg208)II; gjEx866[flp-6::YC3*.*60 elt-2::GFP]*, 6x outcrossed

GJ1498 *tph-1(mg208)II; gjEx523[sra-6::YC3*.*60 elt-2::GFP]*, 6x outcrossed

GJ1499 *odr-3(n1605)V; gjEx523[sra-6::YC3*.*60 elt-2::GFP]*, 6x outcrossed

GJ2202 *gjEx866[flp-6::YC3*.*60 elt-2::GFP]*

GJ2208 *unc-13(e51)I; gjEx866[flp-6::YC3*.*60 elt-2::GFP]*, 1x outcrossed

GJ2209 *unc-13(e51)I; gjEx523[sra-6::YC3*.*60 elt-2::GFP]*, 1x outcrossed

GJ2210 *egl-3(ok979)V; gjEx866[flp-6::YC3*.*60 elt-2::GFP]*, 2x outcrossed

GJ2211 *egl-3(ok979)V; gjEx523[sra-6::YC3*.*60 elt-2::GFP]*, 2x outcrossed

GJ2213 *cat-2(tm2261)II; gjEx523[sra-6::YC3*.*60 elt-2::GFP]*, 1x outcrossed

GJ2214 *cat-2(tm2261)II; gjEx866[flp-6::YC3*.*60 elt-2::GFP]*, 1x outcrossed

GJ2218 *unc-31(e928)IV; gjEx523[sra-6::YC3*.*60 elt-2::GFP]*, 1x outcrossed

GJ2219 *unc-31(e928)IV; gjEx866[flp-6::YC3*.*60 elt-2::GFP]*, 1x outcrossed

GJ2223 *eat-4(ad819)III; gjEx866[flp-6::YC3*.*60 elt-2::GFP]*, 3x outcrossed

GJ2224 *eat-4(ad819)III; gjEx523[sra-6::YC3*.*60 elt-2::GFP]*, 3x outcrossed

GJ2277 *otIs204[ceh-32p::lsy-6 elt-2::GFP]; gjEx866[flp-6::YC3*.*60 elt-2::GFP]I, 1x outcrossed* OH8585 *otIs4 [gcy-7p::gfp]; otEx3822 [ceh-36p::CZ-caspase3(p17) gcy-7p::caspase3(p12)-NZ myo-3p::mCherry]* (ref. 24)

OH8593 *ntIs1 [gcy−5p::GFP lin-15(+)]; otEx3830 [ceh-36p::CZ-caspase3(p17) gcy−5p::caspase3(p12)-NZ myo-3p::mCherry]* (ref. 24)

Germline transformation was performed as described, using an *elt-2p::GFP* construct as co-injection marker^51^. Promoters used for expressing the Yellow Cameleon (YC3.60)^21,22^ construct were *sra-6* for ASH and *flp-6* for ASE.

### Cameleon imaging

Images were acquired with a Zeiss Axiovert 200M microscope, fitted with a Harvard apparatus MC-27 flow chamber. The naïve wash buffer contained 5 mM K_2_HPO_4_/KH_2_PO_4_, pH 6.6, 1 mM MgSO_4_, 1 mM CaCl_2_, the pre-exposure and stimulus buffers contained additional NaCl. The osmolarity of these buffers was set to 325 mosmol, using glycerol, except when NaCl concentrations were too high. Animals were glued onto 2% agarose pads using Nexaband^®^ veterinary glue (World Precision Instruments, Sarasota, Florida). Stimuli were applied by moving a capillary into the buffer close to the nose of the worm. We used a custom automation in Improvision Openlab to control the movement of the capillary and to acquire the images. The acquired image was split into a CFP and YFP part with an Optical Insights Dualview beamsplitter (dichroic mirror 505 nm, 465/30 nm and 535/30 nm emission filters), and the intensities of the CFP and YFP fluorescent areas were recorded, normalized to the 2 seconds prior to the stimulus. The fluorescent ratio was determined by (YFP intensity)/(CFP intensity) −0.6, where the 0.6 factor corrects the bleedthrough of CFP into the YFP channel.

### Behavioral experiments

The response to 25 mM NaCl, with or without pre-exposure to 100 mM NaCl, was assessed as described before^6,42^. Briefly, animals were synchronized by bleaching and grown for 66-72 at 25°C. The animals were washed for 15 minutes with CTX buffer (K_2_HPO_4_/KH_2_PO_4_, pH 6.6, 1 mM MgSO_4_, 1 mM CaCl_2_) with or without 100 mM NaCl and (in a minimal volume) approximately 100 animals were transferred to the center of a quadrant chemotaxis assay plate (Falcon X plate). 2 Quadrants of the assay plate contained CTX agar (1.7% bacto agar, CTX buffer) with 25 mM NaCl, 2 quadrants contained CTX agar without NaCl; 15 minutes before the assay the quadrants were connected by a thin layer of CTX agar. Assay duration was 10 minutes except in the experiments presented in Fig. 8, where animals were followed for 60 minutes. A chemotaxis index was calculated: (*A − C*)/(*A + C*), where *A* is the number of animals at the quadrants with NaCl, and *C* is the number of animals at the quadrants without attractant. Assays were performed in duplicate, at least on two different days. The behavior of animals was always compared with controls performed on the same day(s). Animals that did not move away from the center or were located above the plastic edges were censored.

### Statistical analysis

All experimental results are given as a mean +/-standard error of the mean (SEM). Statistical significance was determined using an ANOVA, followed by a Bonferoni post hoc test.

### Computational modeling

Virtual worms were simulated in the quadrant assay^6,39^ (**Fig. M5** in Supplementary methods) and in the spots assay (**Fig. 9, Fig. M6** in Supplementary methods). In the quadrant assay, every data point was run with 1,000 animals; in the spot assay, we used 500. Quadrant assay simulations were initialized with worms at the center of a plate with Cartesian quadrants of alternating salt concentration. The interface between quadrants was modeled smoothly with a peak concentration gradient of 100 M/m (see Supplementary methods). Simulated behavior was quantified by the chemotaxis index as in the behavioral assay. The spot assay consisted of a hexagonal grid, defined by the spot radius, peak concentration and spot separation distance. Chemotaxis was quantified by hop frequencies.

Model worms consist of a point worm with three sensory neurons ASEL, ASER, and ASH, a single downstream interneuron controlling the pirouette rate and a simplified steering circuit. ASEL threshold modulation is used to model desensitization. Multiplicative gain modulation in ASER is used to model sensitization. ASH recruitment was modeled as a dynamic switch with rates that depend on the history of the NaCl concentration. Sensitization and desensitization rates were fit to match gustatory plasticity rates from the Ca^2+^ imaging results. All virtual assays were simulated in the absence of food.

Steering was implemented by a half-center oscillator circuit capable of generating undulations as well as steering the worm, consistent with behavioral data and neuronal circuit motifs^38,52^. Model parameters were set so the pattern generation was achieved endogenously, but the same model circuit would support alternative (proprioceptively driven) control mechanisms^3^.

To confirm the validity of the steering model to the quadrant assay, we qualitatively compared the behavior in simulations and experiments. As the strongest steering was observed near the quadrant boundaries, we systematically simulated model animals approaching and crossing the boundary at different angles of attack (**Supplementary Fig. 6**). Simulating the locomotion of simplified model animals with only one, fully sensitized sensory neuron and no pirouettes consistently showed that only ASEL influences the direction of the model trajectories across the sharp quadrant boundary, whereas ASER fails to steer the model animal. In these simulations, the comparatively faster rise time of ASEL (on the time scale of an undulation) is required to effect steering while the decay rate of the rectified ASEL response is immaterial. Finally, in our model, the rectification of ASEL, which limits steering to motion up the NaCl gradient is required to avoid negative chemotaxis when heading down the gradient.

Instantaneous changes of bearing to a new random direction were used to mimic a pirouette^3^. The pirouette rate was set by a single neuron, whose activation was up-or down-regulated by incoming negative (aversive) or positive (attractive) sensory signals. The spot assay base pirouette rate was set to 2.1 turns/min^26^, whereas in the quadrant assay, a lower base pirouette rate was used to match experimental trajectories of naïve animals which predominantly used steering to orient themselves (0.3 turns/min in the quadrant assay, n=14 animals).

Sensitization of ASER and ASH can lead to a competition between attractive and aversive responses to salt. To impose the aversive response, we set a much stronger ASH weight onto the pirouette interneuron than that from ASER, thus ‘drowning’ the attractive drive. An alternative ‘blocking’ mechanism whereby ASH actively disrupts signaling along the ASE sensorimotor pathway is equally tenable. In fact, Oda *et al*. showed a complete loss of activity to NaCl downsteps in the AIB interneurons (postsynaptic to ASER) after pre-exposure to NaCl in the absence of food^25^.

To validate our models, we confirmed that model animals exhibited similar motor behavior to those of animals in the quadrant assay (**Fig. M4** in Supplementary methods). Further modeling details are given in Supplementary methods.

## Acknowledgements

We thank R. Tsien, A. Miyawaki and B. Schafer for constructs, B. Schafer and H. Suzuki for help in setting up imaging and for sharing unpublished results, WormBase for providing annotated data on prior work, and the *Caenorhabditis* Genetics Center for strains. We also thank C. Brittin, R. Gudde, I. Hope, R. Hukema, S. Lockery, S. Rademakers and T. Thiele for suggestions. This work was funded by the Center for Biomedical Genetics, the Royal Netherlands Academy of Sciences, ALW/NWO and the EPSRC via grants EP/J004057/1 and EP/N010523/1.

## Author contributions

M.P.J.D., F.S., T.S., O.U., N.C. and G.J. designed the research and analyzed and interpreted the data. M.P.J.D., O.U. and G.J. performed *C. elegans* genetic, imaging and behavioral experiments. F.S. and T.S. developed computational models. M.P.J.D., F.S., T.S., N.C. and G.J. wrote the manuscript.

