## Supplemental material for "Plasticity in gustatory and nociceptive neurons controls decision making in *C. elegans* salt navigation"

**Supplementary information**

Supplementary Figure 1-8
Supplementary Table 1
Supplementary Methods
Supplementary Movie S1-S30

### Supplementary figures

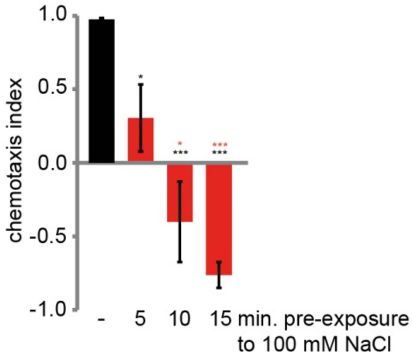


**Supplementary Figure 1. Pre-exposure to NaCl significantly reduces chemotaxis to NaCl and is time dependent.** Wild type animals were tested for chemotaxis to 25 mM NaCl after exposure to 100 mM NaCl for 5-15 min. (red bars), or as a control to a buffer containing no NaCl, for 15 min. (black bar; 5 or 10 min. wash in a buffer without NaCl gave similar results as 15 min. wash). 5-15 min. pre-exposure resulted in a significant decrease of chemotaxis to NaCl (black * $p<0.05$, *** $p<0.001$, compared to no NaCl). 10 or 15 min. pre-exposure resulted in a significantly stronger reduction of chemotaxis (red * $p<0.05$, *** $p<0.001$, compared to 5 min. pre-exposure). 15 min. pre-exposure did not significantly reduce chemotaxis further, compared to 10 min. pre-exposure ($p>0.05$).

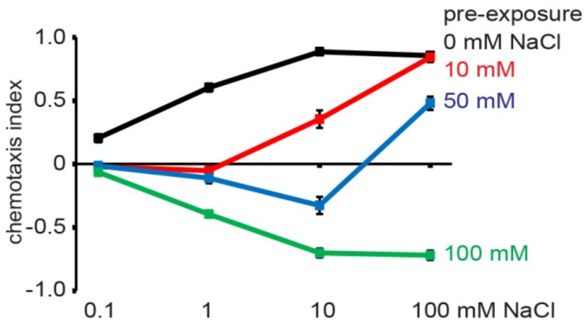


**Supplementary Figure 2. Pre-exposure to NaCl does not affect chemotaxis to higher NaCl concentrations.** Wild type animals were tested for chemotaxis to 0.1, 1, 10 or 100 mM NaCl after exposure to 0, 10, 50 or 100 mM NaCl for 15 min. Pre-exposure strongly affected chemotaxis to NaCl concentrations lower or similar to the pre-exposure concentration but did not affect chemotaxis to higher concentrations: chemotaxis to 10 mM was significantly different from that to 100 mM for animals pre-exposed to 10 mM NaCl ($p<0.001$) or to 50 mM ($p<0.001$), but not after pre-exposure to 0 or 100 mM NaCl ($p>0.05$). Indicated are the averages of 18 assays ± SEM.

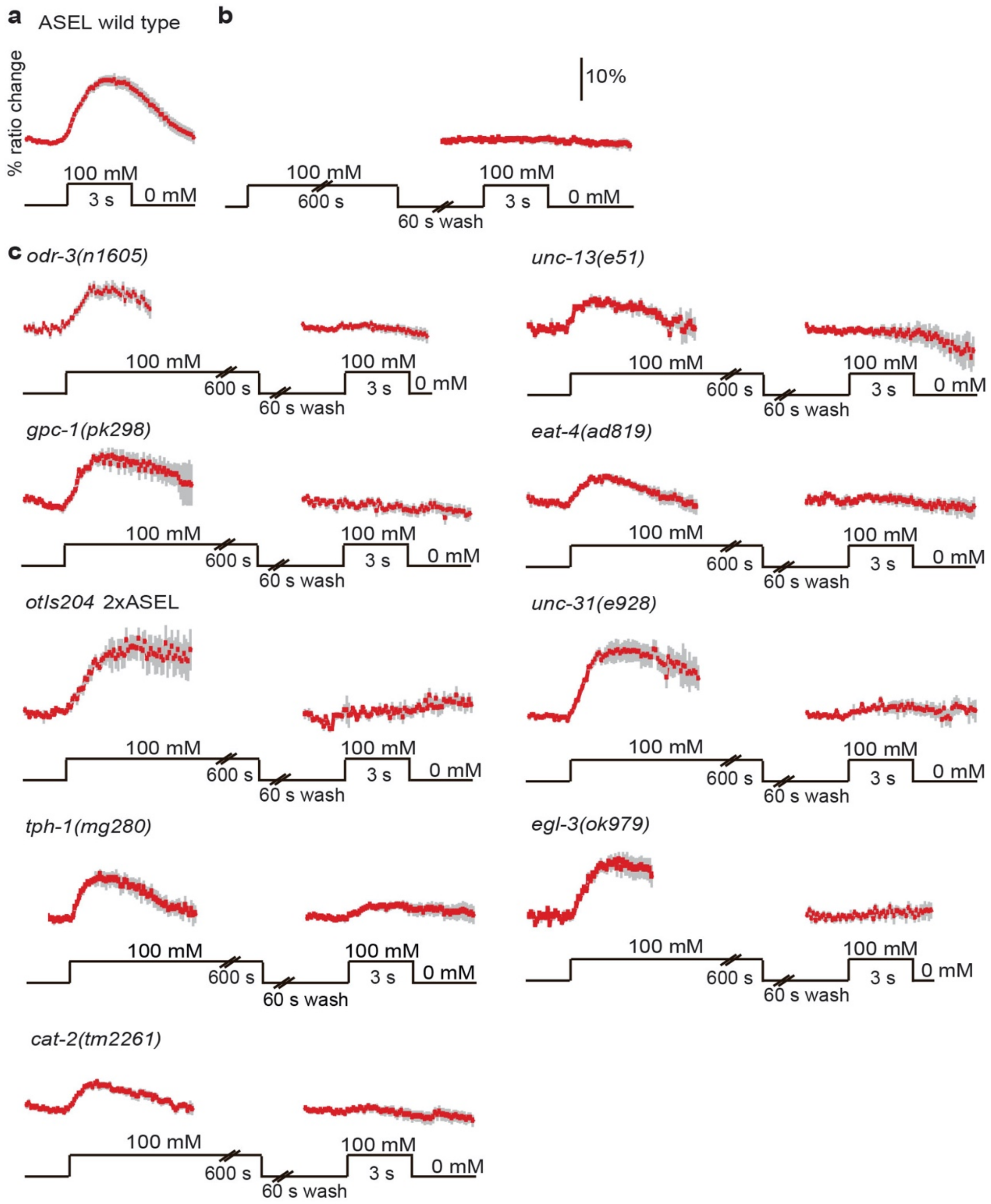


**Supplementary Figure 3. Desensitization of ASEL in various mutant strains.** (**a**) Average $Ca^{2+}$ transients (± SEM) of naïve wild type animals in response to a 3 seconds exposure to 100 mM NaCl (data from Figure 1). (**b**) Average $Ca^{2+}$ transients (± SEM) of wild type animals in response to 100 mM NaCl after 10 minutes exposure to 100 mM NaCl (data from Figure 3). (**c**) Average $Ca^{2+}$ transients (± SEM) in ASEL of *odr-3(n1605)*, *gpc-1(pk298), otIs204*, *tph-1(mg208)*, *cat-2(tm2261)*, *unc-13(e51)*, *eat-4(ad819)*, *unc-31(e928)* and *egl-3(ok979)* animals in response to 100 mM NaCl, either naïvely or after 10 minutes exposure to 100 mM NaCl. The average maximum ratio changes of none of the mutants was significantly different from that of wild type animals (p>0.05). *odr-3(n1605)*: n=7, *gpc-1(pk298)*: n=13, *otIs204*: n=4, *tph-1(mg208)*: n=9, *cat-2(tm2261)*: n=9, *unc-13(e51)*: n=7, *eat-4(ad819)*: n=8, *unc-31(e928)*: n=8 and *egl-3(ok979)*: n=7.

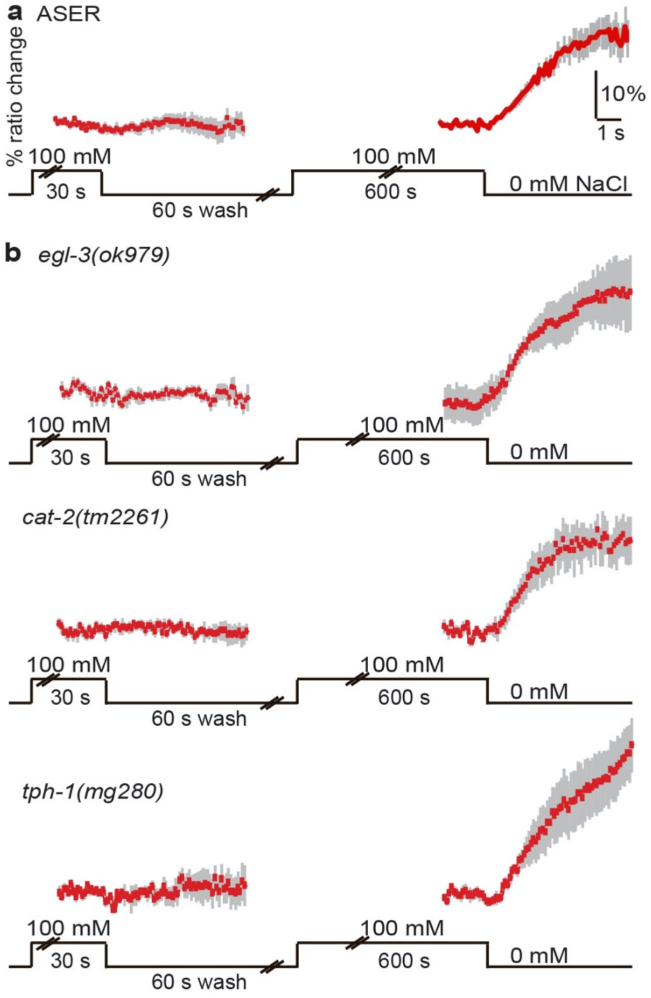


**Supplementary Figure 4. Sensitization of ASER does not involve neuropeptides, dopamine or serotonin.** (**a**) Average $Ca^{2+}$ transients (± SEM) in ASER in wild type animals in response to a decrease in NaCl concentration from 100 mM to 0 mM after 30 or 600 seconds exposure (data from Figure 5). (**b**) Average $Ca^{2+}$ transients (± SEM) in ASER neurons of *egl-3(ok979)*, *cat-2(tm2261)* and *tph-1(mg280)* animals in response to a decrease in NaCl concentration from 100 mM to 0 mM after 30 or 600 seconds exposure. 30 seconds exposure to 100 mM NaCl did not result in a response in ASER, but 10 minutes exposure did result in a $Ca^{2+}$ response. *egl-3(ok979)*: n=4, *cat-2(tm2261)* n=4, *tph-1(mg280)* n=5. The average maximum ratio changes of none of the mutants was significantly different from that of wild type animals ($p>0.05$).

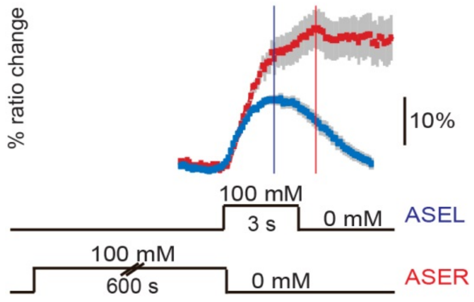


**Supplementary Figure 5. ASEL $Ca^{2+}$ response is faster than ASER response.** Superimposed average $Ca^{2+}$ transients of fully sensitized ASEL (blue) and ASER (red) neurons of animals exposed to 100 mM increases or decreases (600s pre-exposed), taken from Fig. 1 and 2 respectively. Vertical lines denote time of peak depolarization.

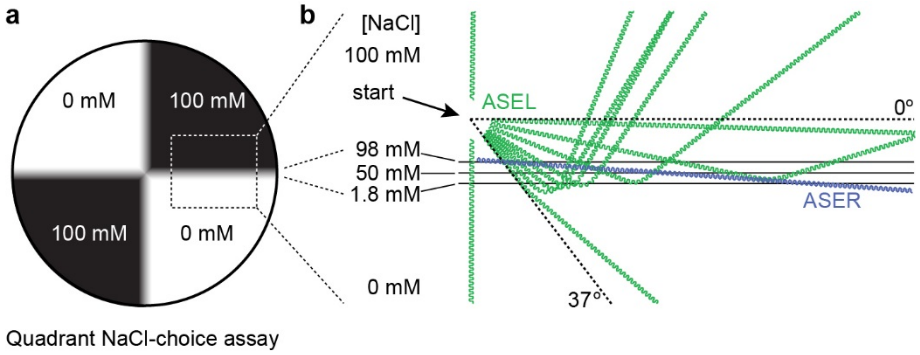


**Supplementary Figure 6. Model worm trajectories on quadrant NaCl-choice assay.** (**a**) Schematic of quadrant NaCl-choice assay[9]. Animals are delivered in the middle of a plate divided in 4 quadrants filled with buffered agar, 2 quadrants with e.g. 100 mM NaCl, 2 quadrants without NaCl, separated by plastic spacers. Just before the assay the area between the quadrants is filled with agar, resulting in a very steep gradient from 100 to 0 mM NaCl (see Supplementary methods and Fig. M5). (**b**) Sample trajectories of virtual worms with either virtual ablations of ASER (ASEL only, in green) or ASEL (ASER only, in blue), initiated (start) in the vicinity of a steep salt transition between 100 and 0 mM NaCl (modeled after the quadrant assay). NaCl concentrations in the transition area have been indicated. For sufficiently shallow angles of attack (down the salt gradient), ASEL successfully mediates strong steering up the salt gradient. The effect of ASER on the steering is negligible. Simulations with virtual ASEL ablations yielded a maximum change of orientation below 2° and only in trajectories close to perpendicular to the salt gradient.

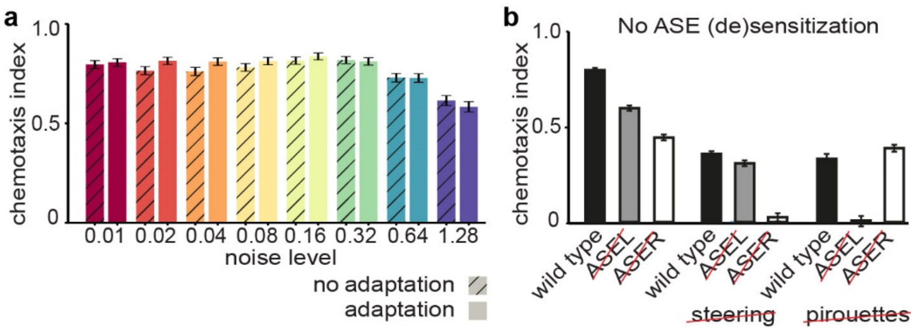


**Supplementary Figure 7. Robustness of the model performance in the virtual quadrant assay is not reduced by ASE (de)sensitization. (a)** Populations of animals display similar CI distributions with and without ASE (de)sensitization. The CI (mean and standard deviation) for 'noisy' populations of $n = 500$ animals. In each animal within each population, ASE parameters were perturbed with the specified noise amplitude (see Supplementary Methods). Performance was measured by the CI of the population after 10 minutes in the quadrant assay. CI levels remained steady with noise amplitudes up to $\sigma$ =0.32. Regardless of CI performance, model worms with (test group) (de)sensitization performed at least as well as those without sensory adaptation (control group). For $\sigma$ >0.32 the performance dropped in both test and control groups. **(b)** ASE (de)sensitization does not contribute to the asymmetric functions of the ASE neurons in our computational model. Chemotaxis index in the virtual quadrant assay of animals with fully sensitized ASEL and ASER. Ablating ASEL or ASER and disabling steering or pirouettes have very similar effects as in animals with wild type ASE (de)sensitization (Fig. 8c).

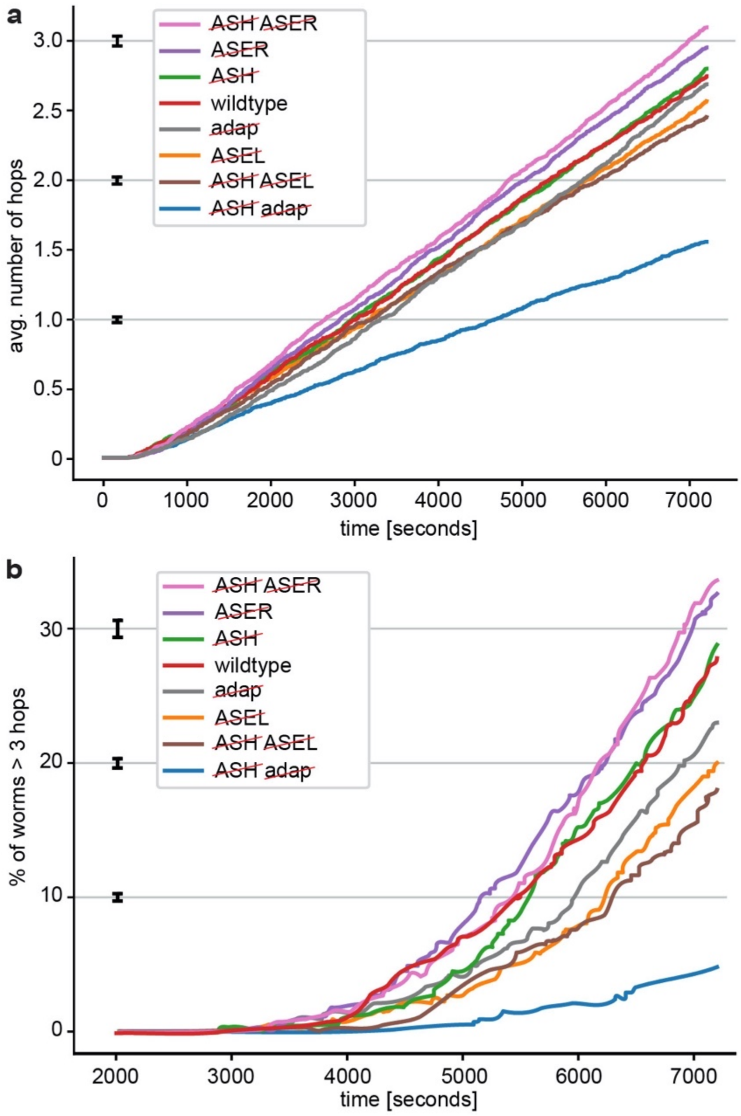


**Supplementary Figure 8. Exploration behavior over time in the spot assay.** The time evolution of the hop counts from Fig. 10. A hop from one spot to another is registered once a worm trajectory meets a spot for the first time, after leaving another spot. (**a**) The average number of hops per worm over time, of wild type worms and worms in which specific cells were ablated (e.g. ~~ASER~~) or in which ASEL and ASER are always fully sensitized (~~adap~~). The average number of hops is hardly affected by ablating ASH or ASEL at all times. (**b**) Fraction of worms that hopped at least three times. The variance of the hop counts vastly differs, leading to different numbers of worms reaching 3 hops within the time window (Fig. 10d). This indicates that the overall exploration success of the worms is robust.

**Supplementary Table 1. ASEL, ASER and ASH $Ca^{2+}$ responses**

| Neuron | Figure | Pre-exp. time (s) | Wash time (s) | Exposure time (s) | Exposure [NaCl] (mM) | n | Response Responding | Response Index[1] | Rise time (s) Mean ± SD | Rise time (s) Range |
|---|---|---|---|---|---|---|---|---|---|---|
| ASEL | 1a,b | | | 3 | 10 | 17 | 10 | 0.59 | 1.6 ± 0.6 | 0.5 - 2.8 |
| | | | | 3 | 100 | 42 | 37 | 0.88 | 1.9 ± 0.7 | 0.7 - 3.8 |
| | | | | 3 | 200 | 16 | 9 | 0.56 | 1.9 ± 0.7 | 1.0 - 3.6 |
| | | | | 3 | 300 | 17 | 15 | 0.88 | 2.0 ± 0.7 | 1.0 - 3.4 |
| | | | | 3 | 500 | 11 | 8 | 0.73 | 1.5 ± 0.6 | 0.5 - 2.3 |
| ASEL | 3a,b | 60 | 60 | 3 | 100 | 13 | 10 | 0.77 | 2.1 ± 0.9 | 1.4 - 4.2 |
| | | 120 | 60 | 3 | 100 | 9 | 8 | 0.89 | 2.3 ± 0.9 | 1.0 - 3.8 |
| | | 300 | 60 | 3 | 100 | 8 | 4 | 0.50 | 1.4 ± 1.2 | 0.4 - 3.3 |
| | | 600 | 60 | 3 | 100 | 16 | 0 | 0.00 | | |
| ASEL | 3e,f | 600 | 120 | 3 | 100 | 8 | 6 | 0.75 | 3.0 ± 1.2 | 1.5 - 4.8 |
| | | 600 | 300 | 3 | 100 | 6 | 5 | 0.83 | 2.9 ± 1.3 | 1.1 - 4.9 |
| ASH | 1d,e | | | 3 | 100 | 18 | 5 | 0.28 | 2.7 ± 0.7 | 1.8 - 3.4 |
| | | | | 3 | 200 | 18 | 5 | 0.28 | 1.7 ± 0.7 | 0.7 - 2.6 |
| | | | | 3 | 300 | 22 | 16 | 0.73 | 2.4 ± 0.6 | 1.7 - 4.2 |
| | | | | 3 | 500 | 19 | 16 | 0.84 | 2.3 ± 0.6 | 1.5 - 3.6 |
| ASH | 4a,b | 600 | | 3 | 200 | 17 | 16 | 0.94 | 2.6 ± 0.4 | 1.8 - 3.5 |
| | | 600 | | 3 | 300 | 12 | 10 | 0.83 | 3.0 ± 0.3 | 2.6 - 3.6 |
| | | 600 | | 3 | 500 | 24 | 19 | 0.79 | 2.9 ± 0.7 | 2.2 - 4.6 |
| ASER | 1c | | | 3 | 10 | 8 | 0 | 0 | | |
| | | | | 3 | 100 | 30 | 0 | 0 | | |
| | | | | 3 | 200 | 5 | 0 | 0 | | |
| | | | | 3 | 300 | 9 | 0 | 0 | | |
| | | | | 3 | 500 | 4 | 0 | 0 | | |
| ASER | 2a,b | | | 30 | 100 | 8 | 1 | 0.13 | 8.3 | 8.3 |
| | | | | 60 | 100 | 15 | 10 | 0.66 | 6.0 ± 4.3 | 0.4 - 13.0 |
| | | | | 120 | 100 | 4 | 4 | 1.00 | 5.5 ± 3.0 | 2.0 - 8.5 |
| | | | | 300 | 100 | 5 | 3 | 0.60 | 3.4 ± 0.9 | 2.2 - 4.4 |
| | | | | 600 | 100 | 29 | 25 | 0.86 | 6.2 ± 3.4 | 1.8 - 13.5 |

[1]. The fraction of animals responding has been indicated as the response index (# animals responding / # animals tested (n)).

### Supplementary methods

#### Computational model

A computational simulation framework was developed to study *in silico* the behavior of single model worms confronted with the quadrant assay. A variation of this computational model has previously been used to model a *C. elegans* decision making assay[1]. Our model of the animal assumes that during locomotion, the body follows the head, allowing us to focus on sensory-motor control of a point worm. Thus, at each point in time, animals are modeled as a point $(x(t), y(t))$ representing the head heading towards an angle, $\theta(t)$. The head moves at a fixed speed, $v = 0.22$ mm/s, resulting in an overall body speed of $v = 0.1$ mm/s. Our model *C. elegans* sense their environment using a simplified nervous system which dynamically controls the direction of locomotion[1,2].

To study gustatory plasticity and the effects of salt adaptation on navigation, our model circuit contained three sensory neurons (ASEL, ASER and ASH) that synapse directly onto a simplified motor system (**Fig. M1**). The motor output consists of undulations that are modulated by the sensory system to generate steering, in addition to instant turning events representing pirouettes. All neuronal equations and parameters, including the sensory responses and motor neuron dynamics were constrained by $Ca^{2+}$ imaging data[3,4] and published behavioral data[4-8].

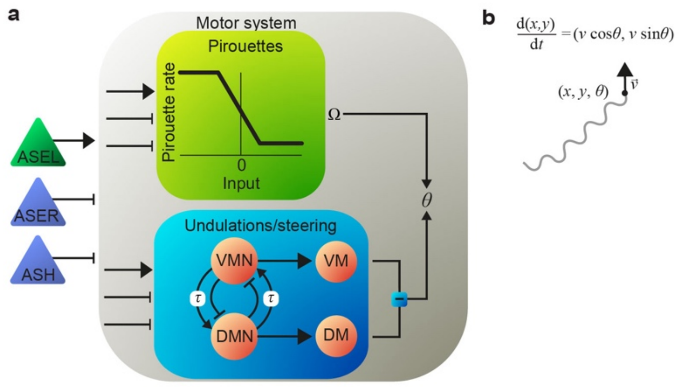


**Fig. M1.** Schematic of the computational model. (**a**) The model circuit contains three sensory neurons, ASEL, ASER and ASH (triangles), and a motor circuit (grey box). Each sensory neuron projects onto all three components of the motor circuit (Pirouettes, VMN, DMN). A motor command unit (green box) regulates the pirouette rate as a function of sensory input. Positive synaptic input suppresses the probability of a pirouette; negative input enhances it. A pirouette ($\Omega$) is modeled as an instantaneous, random reorientation. A half-center oscillator circuit (blue box) generates undulations and steers the worm in response to synaptic inputs from sensory neurons: Two motor neurons (VMN, DMN) drive model muscles (VM, DM) on the respective sides of the body. Each delay unit $\tau$ and muscle (VM, DM) in the motor circuit is modeled as an implicit neuronal unit. The output of the sensory motor circuit is an instantaneous orientation, $\theta(t)$. (**b**) Visualization of the worm coordinates in time. An animal at time t is represented by a point, $(x(t), y(t))$, in a two-dimensional space that moves with fixed speed, $v$, in a direction, $\theta(t)$.

We followed an incremental approach for parameter tuning of the model. First, sensory neuron rate parameters $(\alpha_i, \beta_i, \gamma_i)$ were manually fit to match $Ca^{2+}$ imaging traces of naïve animals (**Fig. M3**). In **Figure M2**, $\alpha_i$ determines the steepness of the fast conductance; $\gamma_i$ determines the steepness of the slow conductance; and $\alpha_i/\beta_i$ determines the saturation amplitude of F (in arbitrary units). Together, these parameters determine the current profile $(\mathrm{F} - \mathrm{S})$. The current-voltage relation (Equation M1 below) was not fit (parameter values used were the same as in as Ghosh *et al.* (2016)[1]).

The motor circuit (synaptic weight parameter set in **Table M1**) was independently modeled to obtain realistic undulations and pirouette rates. Modeling salt navigation behaviors only required setting the synaptic weights from salt sensory neurons onto the motor circuit (to match observed steering and turning rates in the quadrant assay). This modeling approach ensures that the model is maximally constrained and should enhance its predictive potential. Once the model was fully parametrized, it was tested, first on the quadrant assay, and second, on a new *in silico* spot assay. Only the base pirouette rate was adjusted in the spot assay, in agreement with observations on related assays[8].

**Model neurons**

We modelled all neurons as leaky integrators (with arbitrary units)

$$\tau_{\mathrm{m}} \frac{\mathrm{d}V_i}{\mathrm{d}t} = -V_i + V_{0,i} + \sigma(I), \qquad \text{(M1)}$$

where $V_i$ is a voltage like variable for neuron $i$, also referred to here as neuronal activation; $\tau_{\mathrm{m}}$ is a neuronal time constant; finally, $V_{0,i}$ represents the resting potential (set throughout to zero). The function $\sigma(I)$ is a sigmoid defined as

$$\sigma(I) = \tanh(bI), \qquad \text{(M2)}$$

where $b$ is a gain parameter and the input current is denoted $I = I(t)$ (with the implicit neuronal index $i$, **Table M1**). Thus, a neuron's activation ranges from -1 to 1 and converges to $\sigma(I)$ with a timescale of $\tau_{\mathrm{m}}$ for fixed input, $I$. This formulation is borrowed from rate neuron models and naturally exhibits thresholding and saturation of activation, as observed in *C. elegans* sensory neurons[3,4,9]. As the bilateral ASHL/ASHR neuronal pair is coupled by gap junctions[10] and ASH Left and Right neurons are known to respond identically[4], the ASH neuron pair was collapsed into a single model neuron.

**Table M1.** Neuronal parameters and synaptic weights.

| Neuronal parameters | Value | Description |
|---|---|---|
| $\tau_{\mathrm{m}}$ | 0.5 s | Neuronal time constant |
| $V_{0,i}$ | 0 | Resting potential |
| $b$ | 2 | Neuronal gain |
| **Synaptic weights** | **Value** | **Description** |
| $W_{\mathrm{ASEL,M}}$ | 0.5 | ASEL onto DMN and VMN |
| $W_{\mathrm{ASER,M}}$ | -0.5 | ASER onto DMN and VMN |
| $W_{\mathrm{ASH,M}}$ | 0.5 | ASH onto DMN and VMN |
| $W_{\mathrm{ASEL,\Omega}}$ | -0.25 | ASEL onto Ω |
| $W_{\mathrm{ASER,\Omega}}$ | -0.25 | ASER onto Ω |
| $W_{\mathrm{ASH,\Omega}}$ | 1 | ASH onto Ω |
| $W^{+}_{\mathrm{D,V}}$ | 0.88 | DMN to VMN excitation (to, from hidden neuron) |
| $W^{+}_{\mathrm{V,D}}$ | 0.88 | VMN to DMN excitation (to, from hidden neuron) |
| $W^{-}_{\mathrm{D,V}}$ | -1.4 | DMN to VMN inhibition |
| $W^{-}_{\mathrm{V,D}}$ | -1.4 | VMN to DMN inhibition |

The input current *I* sums over all synaptic and sensory contributions, $I = I_{\text{syn}}(t) + I_{\text{sens}}(t)$, where, assuming graded synaptic transmission, $I_{\text{syn}} = \sum_j W_{ji} V_j$, is a weighted sum over all presynaptic neuron activations and $I_{\text{sens}}$ denotes the stimulus response current in sensory neurons (given in Equations (M5)–(M7)). In line with the unrectified nature of graded synapses in *C. elegans*[11], hyperpolarizing a neuron will effectively reverse the polarity of its efferent synaptic transmission (with hyperpolarizing and depolarizing postsynaptic transmission across excitatory and inhibitory synapses, respectively).

**Sensory neurons**

Our model focuses on sensory responses to NaCl. In this study we looked exclusively at ASEL, ASER and the ASH sensory neuron pair. $Ca^{2+}$ imaging data has shown that multiple sensory neurons in *C. elegans* respond to a change in a stimulus, rather than to the stimulus intensity directly. The response often approximates a time-filtered time derivative over the stimulus input[3,4,12,13]. Moreover, in many cases, changes in the stimulus level selectively produce a rectified depolarizing response with a characteristic rise time, followed by a slower decay to rest. In particular, ASEL and ASH neurons respond with a transient depolarization to increases in NaCl concentration (in the case of ASH, only to high concentrations above ~200 mM indicating a response to high osmolarity, rather than NaCl *per se*)[3]. In addition, ASER and ASH show a transient depolarization in response to decreases in the NaCl concentration[4].

The biphasic nature of the derivative-like responses seen in *C. elegans* sensory neurons suggests a model in which sensory neurons use two driving forces or conductances with opposite effect and a separation of timescales[13]. Importantly, the characteristic rise and decay times may be neuron specific (and possibly stimulus specific). Although detailed electrophysiological characterization is not yet available for these neurons, $Ca^{2+}$ imaging with a variety of indicators[3,4,14], including our own data here, indicates that the ASER response (in particular the associated decay time) to a decrease in NaCl is significantly slower than the ASEL response to a NaCl increase.

In our model, the sensory contribution to the neuronal activity is denoted by a current, $I_{\text{sens}}$. We modeled sensory responses using a fast component, $F$, and a delayed rectifying component, $S$ (**Fig. M2a**). The sensory current $I_{\text{sens}}$ is given by the difference between the fast and slow components, subject to either positive rectification (in ASEL), or reversal (ASER) depending on the cell. We represent this additional transformation generically as

$$I_{\text{sens}} = f_i(F - S) \qquad \text{(M5)}$$

The function, $f_{\text{ASEL}}(x) = \max(0, x)$, enables ASEL to respond only to NaCl concentration increases; $f_{\text{ASER}}(x) = -x$ allows ASER to respond to NaCl concentration decreases. The absence of rectification in ASER captures the effect of hyperpolarization in response to concentration increases. ASH is modeled without rectification or reversal, $f_{\text{ASH}}(x) = x$. Fast and slow components evolve according to

$$\frac{\mathrm{d}F}{\mathrm{d}t} = D_i \alpha_i \max(0, C_{\log} - C_0) - \beta_i F, \qquad \text{(M6)}$$

$$\frac{\mathrm{d}S}{\mathrm{d}t} = \gamma_i (F - S). \qquad \text{(M7)}$$

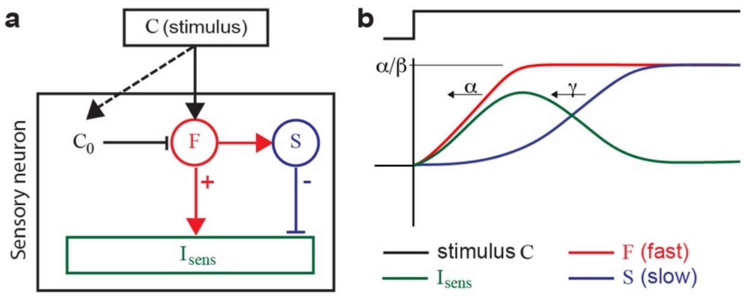


**Fig. M2. Diagram of a sensory neuron and its response.** (**a**) A stimulus $C$ evokes a rapid response, $F$, which in turn drives a slow, delayed response, $S$. The response $F$ is subject to a threshold, $C_0$, depicted here as a bias input. The current $I_{\mathrm{sens}}$ is a function over the difference, $F - S$, such that $S$ acts as a delayed rectifier. In ASEL, $C_0 \geq 0$ is a slowly adapting baseline that desensitizes the sensory neuron to the ambient stimulus strength (dashed arrow). (**b**) Schematic traces illustrating the response $I_{\mathrm{sens}}$ to a stimulus, $C$, given by the difference, $F - S$, (where $C_0 = 0$). $\alpha_i$ determines the steepness of the fast conductance; $\gamma_i$ determines the steepness of the slow conductance; and $\alpha_i/\beta_i$ determines the saturation amplitude of F (in arbitrary units).

The fast component, $F$, responds to changes in the NaCl concentration with a characteristic rate $\beta_i$. The slow component, $S,$ follows $F$ with a delay, producing the transient response (**Fig. M2b**). We used $Ca^{2+}$ imaging traces under fully sensitized conditions to calibrate the global timescale $\tau_{\mathrm{m}}$ (in compliance with the Motor circuit, **Fig. M1**), as well as to fit the timing coefficients $\alpha, \beta$ and $\gamma$ for the ASEL response (**Fig. M3a**).

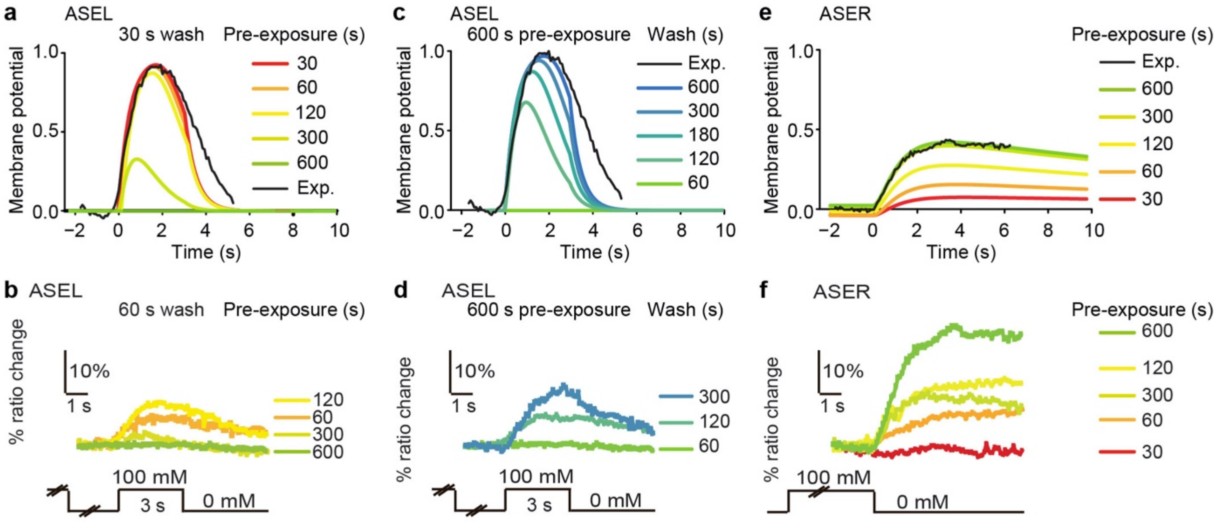


**Fig. M3. Model sensory neuron responses.** (**a**) Model ASEL responses to 3 seconds exposure to 100 mM NaCl (starting at 0 seconds) after pre-exposure of varying durations to 100 mM NaCl and a 30 seconds wash. A representative $Ca^{2+}$ imaging trace (30 seconds pre-exposure to 100 mM NaCl, 30 seconds wash) in black, with vertical scaling to the amplitude of the corresponding model trace. (**b**) *In vivo* counterpart of (a): average $Ca^{2+}$ transients in ASEL in response to 100 mM NaCl after 60-600 seconds pre-exposure and 60 seconds wash. Data from Fig. 3a. (**c**) ASEL responses to 3 seconds exposure to 100 mM NaCl after 600 seconds pre-exposure to 100 mM NaCl and a wash of varying durations. (**d**) *In vivo* counterpart of (c): average $Ca^{2+}$ transients in ASEL in response to 100 mM NaCl after 600 seconds pre-exposure and 60-300 seconds wash. Data from Fig. 3a,e. (**e**) ASER responses to a decrease in NaCl from 100 to 0 mM (at 0 seconds) after pre-exposure to 100 mM NaCl of varying durations. A representative $Ca^{2+}$ imaging trace (600 seconds pre-exposure to 100 mM NaCl) in black, with vertical scaling to the amplitude of the corresponding model trace. (**f**) *In vivo* counterpart of (e): average $Ca^{2+}$ transients in ASER after 30-600 seconds pre-exposure to 100 mM NaCl. Data from Fig. 2a.

The order of magnitude for gain factor $b$ was chosen so the response is clipped within the range of input current variations induced by pre-exposure (see below). We used $b$ to tune the shape of the head trajectory (**Fig. M4a**). For parsimony, we used the same gain factor and neuron timescale in all motor and sensory neurons. The timing coefficients for ASER where chosen similarly, to match the $Ca^{2+}$ imaging above (**Fig. M3c**). Parameter values are listed in **Table M2**.

**Table M2.** Sensory neuron parameters.

| Sensory neuron parameters | Value | Description |
|---|---|---|
| $\alpha_{\mathrm{ASEL}}$ | 50 $s^{-1}$ | ASEL depolarization rate |
| $\beta_{\mathrm{ASEL}}$ | 8 $s^{-1}$ | ASEL leak rate |
| $\gamma_{\mathrm{ASEL}}$ | 1 $s^{-1}$ | ASEL rectification (repolarization) rate |
| $\delta_{\mathrm{ASEL}}^{\mathrm{ON}}$ | 0.006 $s^{-1}$ | ASEL desensitization rate |
| $\delta_{\mathrm{ASEL}}^{\mathrm{OFF}}$ | 0.004 $s^{-1}$ | ASEL desensitization relaxation rate |
| $D_{\mathrm{ASEL}}$ | 1 | ASEL gain adaptation factor |
| $\alpha_{\mathrm{ASER}}$ | 0.25 $s^{-1}$ | ASER activation rate |
| $\beta_{\mathrm{ASER}}$ | 1 $s^{-1}$ | ASER leak rate |
| $\gamma_{\mathrm{ASER}}$ | 0.05 $s^{-1}$ | ASER rectification rate |
| $\delta_{\mathrm{ASER}}^{\mathrm{ON}}, \delta_{\mathrm{ASER}}^{\mathrm{OFF}}$ | 0 $s^{-1}$ | ASER desensitization rates (off) |
| $\lambda$ | 0.01 $s^{-1}$ | ASER gain adaptation relaxation rate |
| $\alpha_{\mathrm{ASH}}$ | 0.1 $s^{-1}$ | ASH activation rate |
| $\beta_{\mathrm{ASH}}$ | 0.6 $s^{-1}$ | ASH leak rate |
| $\gamma_{\mathrm{ASH}}$ | 0.01 $s^{-1}$ | ASH rectification rate |
| $\delta_{\mathrm{ASH}}^{\mathrm{ON}}, \delta_{\mathrm{ASH}}^{\mathrm{OFF}}$ | 0 $s^{-1}$ | ASH desensitization rates (off) |
| $D_{\mathrm{ASH}}$ | 1 | ASH gain adaptation factor |
| $C_{\mathrm{min}}$ | 1 mM | Unit scaling factor |

In our data (supported by Oda *et al.*[14]) ASEL showed slow desensitization and ASER slow sensitization (main text, **Figs. 1** and **2**). Specifically, ASEL failed to respond to an increase in NaCl concentration after a pre-exposure of ~600 s. However, after such a pre-exposure ASEL did respond to larger increases in NaCl concentration (main text, **Fig. 2c**). This response hints at an adaptive threshold and, together with the wide dynamic range of responses, is reminiscent of Fechner's law, typically expressed as $r \propto \log s$, where $r$ and $s$ denote the response and signal, respectively. The incremental form of the law $\Delta r \propto \frac{\Delta s}{s}$ describes the minimal stimulus change needed to evoke a response, relative to some baseline[15]. Accordingly, in our model, the slow sensory adaptation of ASEL to NaCl is implemented with a slowly evolving concentration sensitivity threshold, $C_0$ (**Fig. M2**, **M3**)

$$\frac{\mathrm{d}C_0}{\mathrm{d}t} = \delta_i^{\mathrm{ON}} C_{\mathrm{log}} - \delta_i^{\mathrm{OFF}} C_0 \ .$$

In addition, the fast component $F$ integrates over the logarithm of the NaCl concentration, capturing the wide dynamic range of behavioral responses to NaCl

$$C_{\mathrm{log}} = \log\left(1 + C_{(x,y,t)}/C_{\mathrm{min}}\right) / \log\left(1 + (C_{\mathrm{max}}/C_{\mathrm{min}})\right),$$

where $C_{\min}$ is a unit scaling factor, that imposes an effective lower cut-off to the sensory dynamic range, grossly representing the minimal concentration that can evoke a behavioral response[16]. Note that in our model, threshold adaptation only applies to ASEL, thus only there $\delta_i^{\mathrm{ON}}$ and $\delta_i^{\mathrm{OFF}}$ are nonzero.

In contrast to the slow desensitization of ASEL, we and Oda *et al.*[14] have shown that ASER *sensitizes* over time: longer NaCl pre-exposure results in a stronger depolarizing response to decreases in NaCl concentration (main text, **Fig. 1**). This sensitization is captured in our model with a multiplicative gain parameter, $D_{\mathrm{ASER}}$: a slowly evolving measure of the presence of NaCl such that the fast component $F$ increases with the pre-exposure duration, as measured experimentally (**Fig. M3**). The gain parameter evolves according to

$$\frac{\mathrm{d}}{\mathrm{d}t} D_{\mathrm{ASER}} = \lambda C_{\log}(1 - D_{\mathrm{ASER}}) - \lambda D_{\mathrm{ASER}}.$$

The initial activities of the neurons are all set to rest (0) at $t = 0$. Further details of the initial conditions are given below.

**ASH recruitment**

Recruitment of the ASH sensory neuron is modeled as a stochastic switch between on and off states, with switching rates that vary with the animal's NaCl exposure history. In its off or *unrecruited* state, ASH is assumed to respond only to dangerous concentrations of NaCl (>300 mM), whereas in its on or *recruited* state, it responds to low concentrations of NaCl as well. Specifically, in the recruited state, the ASH sensory neuron is modeled as identical to ASER, up to a sign reversal.

Let $\rho$ denote the propensity of ASH to be recruited, which varies monotonically from 0 (no NaCl) to 1 (saturated exposure to NaCl). The recruitment rate evolves according to

$$\frac{\mathrm{d}\rho}{\mathrm{d}t} = \kappa\left(\frac{C_{\log}}{C_{\max}} - \rho\right).$$

Hence, $\rho$ converges to the ratio of the current concentration and a 'maximum' concentration with a rate, $\kappa$. Let the transition rates $\alpha_{\mathrm{rec}}$ and $\beta_{\mathrm{rec}}$ denote the recruitment and 'unrecruitment' rates respectively, such that

$$\mathrm{OFF} \underset{\beta_{\mathrm{rec}}}{\overset{\alpha_{\mathrm{rec}}}{\rightleftharpoons}} \mathrm{ON},$$

with $\alpha_{\mathrm{rec}} = \frac{\rho}{\tau_\alpha}$ and $\beta_{\mathrm{rec}} = \frac{1-\rho}{\tau_\beta}$. The steady state probabilities $P^*(\mathrm{ON})$ and $P^*(\mathrm{OFF})$ for occupying the on (recruited) and off (unrecruited) states are then given by

$$P^*(\mathrm{ON}) = \frac{\alpha_{\mathrm{rec}}}{\alpha_{\mathrm{rec}} + \beta_{\mathrm{rec}}}$$

$$P^*(\mathrm{OFF}) = \frac{\beta_{\mathrm{rec}}}{\alpha_{\mathrm{rec}} + \beta_{\mathrm{rec}}}.$$

The above formulation of the recruitment process allows for a separation of time scales which is not needed here, so, for simplicity and parsimony, we simplify it by collapsing all three timescales to one, such that $\tau_\alpha = \tau_\beta = 1/\kappa$. Default parameter values are given in **Table M3**.

We note that ASH recruitment is unlikely to be as simple as an instantaneous and stochastic on-off switch. However, modeling the recruitment kinetics in greater detail would require sufficiently resolved kinetic data (on the fraction of responders as a function of time and concentration). Our data showed a minority of responders to low NaCl concentrations without pre-exposure, and a very high fraction of responders after 600 sec exposure. Without a clear indication of a change in the response amplitude, we opted for a parsimonious binary (on-off) model of ASH responsiveness (to low NaCl concentrations) and our model of the recruitment kinetics aimed only to capture the observed statistics with minimal assumptions. Modeling the recruitment of the ASH sensory neuron as a stochastic switch between on and off states sufficed for capturing the fractions of recruited ASH neurons on salt, but alternative (e.g. continuous) models are possible. Switching rates in this model vary with the animal's NaCl exposure history (a necessary assumption). In animals that have not been pre-exposed to NaCl, ASH is in its ‘off ‘ or unrecruited state. It is implicitly assumed that in its unrecruited state, ASH neurons respond only to dangerous concentrations of NaCl. In animals that were pre-exposed to NaCl, ASH is in its ‘on’ or recruited state, hence responding to low concentrations of NaCl as well.

**Table M3.** ASH recruitment parameters.

| ASH recruitment | Value | Description |
|---|---|---|
| $C_{\mathrm{max}}$ | 100 mM | NaCl saturation level for recruitment $(\rho \to 1)$ |
| $\kappa$ | 0.001 $s^{-1}$ | Common rate parameter for recruitment kinetics |

**Motor system**

Undulations and steering are modeled by a symmetric oscillator motif, consisting of reciprocal inhibitory and delayed excitatory connections (**Fig. M1**). We used *hidden* interneurons to create a delayed connection between motor neurons. Thus, the delayed excitatory connection from VMN to DMN is implemented as two connections, one from VMN to the hidden interneuron, and another from the hidden interneuron to DMN, using the neuronal time constant of the hidden interneuron, $\tau_{\mathrm{m}}$, as a synaptic delay. The reciprocal connection from DMN to VMN is identical. The model contains two separate motor outputs: undulations and pirouettes. Both can be modulated by sensory inputs.

**Undulations**

Two motor neurons (VMN, DMN) in a half-center oscillator like configuration (**Fig. M1**) are capable of generating and maintaining stable oscillations as well as to steer the worm (**Fig. M4**).

The reciprocal connectivity pattern is reminiscent of connectivity found in several classes of head motor neurons in *C. elegans*. Compared to more compact models, e.g. Izquierdo and Lockery[2], our approach allows for the modulation of the undulation frequency as well the amplitude, more closely matching observed trajectories of worms in the choice assay, especially in the vicinity of the quadrant boundaries (**Fig. M4**).

In contrast, with a simpler model relying on a master pacemaker (adapted from Izquierdo and Lockery[2]), we were unable to account for a significant proportion of worm trajectories without changing the undulation amplitudes to unrealistic levels (data not shown).

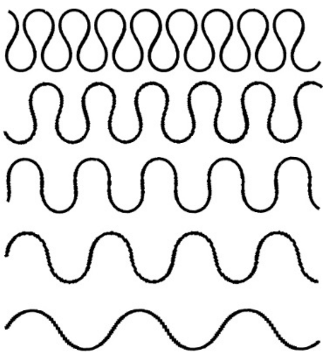


**Fig. M4. Model worm trajectories and motor neuron activities.** The neuronal gain parameter $b$ determines the head undulatory trajectory, with $b = 2$, giving the realistic trajectory. We set the head speed $v$ to obtain an overall speed of 0.1 mm/s resulting in about 23 undulations per cm at a frequency of 0.23 Hz.

In the absence of sensory input, the motor circuit will produce stable oscillations, facilitated by fast reciprocal inhibition that is released by the delayed reciprocal excitation (**Fig. M1**). Any activity in one of the oscillating neurons will cause fast inhibition of the other followed by slower excitation and subsequently inhibition of the originally active neuron. Thus, the frequency and amplitude of the oscillations are determined by the timescales of the neurons, $\tau_{\mathrm{m}}$, the connection strengths of the reciprocal inhibition, and delay in the delayed reciprocal connections. In our model, $\tau_{\mathrm{m}}$, remained fixed, leaving the three pairs of reciprocal connection strengths as parameters to tune the oscillator.

Since the circuit configuration and all parameters are symmetric, symmetry breaking is required to initiate oscillations. Indeed, when VMN and DMN are equally active, the circuit does not oscillate. However, any small difference in activity (or initial conditions) is amplified by the mutual inhibition. In the model, a start 'signal' $V_{\mathrm{start}}$ is given for a short duration $\tau_{\mathrm{start}}$ with opposite polarity to the two motor neurons VMN and DMN, causing them to diverge in activity. Motor output parameter values are given in **Table M4**.

Similar to other point models of *C. elegans*[2,17,18], the direction of locomotion, $\theta$, changes as a function of the difference in activity of the dorsal and ventral motor neurons:

$$\frac{\mathrm{d}\theta}{\mathrm{d}t} = \omega(V_{\mathrm{VMN}} - V_{\mathrm{DMN}}) ,$$

where $\omega$ is the steering strength. Model worms move with constant velocity $v$ along the direction vector $\theta$ according to

$$\frac{\mathrm{d}(x,y)}{\mathrm{d}t} = v(\sin\theta, \cos\theta) .$$

The speed was chosen so the model worm performed realistically on the assay (see **Fig. M5a**).

**Table M4.** Motor output parameters.

| Motor output parameters | Value | Description |
|---|---|---|
| $v$ | 0.22 mm $s^{-1}$ | Locomotion speed |
| $\omega$ | 1.5 $s^{-1}$ | Steering strength/angular speed |
| $V_{\mathrm{start}}$ | 0.001 | Activity of oscillator start signal |
| $\tau_{\mathrm{start}}$ | 0.01 s | Duration of oscillator start signal |
| $\omega_{\Omega}$ | 0.005 $s^{-1}$ | Base pirouette rate |

**Pirouettes**

Instant turning events are executed by resetting the orientation of the movement of the point worm. The probability of a pirouette per unit time $P_\Omega$ is encoded by the activation of the pirouette command unit $V_\Omega$ and given by a smooth and monotonically decreasing function

$P_\Omega(V_\Omega) = w_\Omega \exp(-V_\Omega)$,

where $w_\Omega$ is the base pirouette rate. The parameters were chosen such the base rate is approximately 2.1 pirouettes per minute, and suppression of pirouettes (for sufficiently hyperpolarized values of the neuron) is possible.

When a pirouette is executed, the heading, $\theta$, is instantaneously set to a random angle drawn from a uniform distribution.

**The Model Choice Assay**

Simulations are performed on a virtual two-dimensional 20 cm plate, which animals cannot leave. Upon reaching the edge of the plate, virtual animals are reoriented to a random angle drawn from a uniform distribution. Reorientation can occur multiple times in succession if worms continue to hit the edge. A choice assay (**Fig. M5**) was implemented to reproduce an experimental assay designed to test naive and conditioned responses to NaCl[5]. Supplementary movies S1-S14 show 500 simulated ASH-virtual ablated worms, with/without (de)sensitization in ASEL or ASER, ASEL or ASER-virtual ablated, and/or ablating the synaptic connections to the steering circuit or to the pirouette neuron.

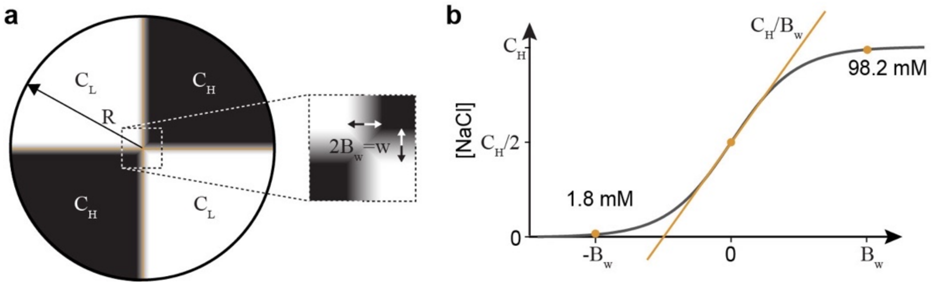


**Fig. M5. The choice assay used in our simulations.** (**a**) A plate of radius $R$ is prepared with alternating quadrants of uniform high ($C_\mathrm{H}$=100 mM, black), and low ($C_\mathrm{L}$=0 mM, white) NaCl concentrations. **(b)** Between quadrants, a sigmoidal concentration profile is imposed such that 96% of the concentration range is spanned within 4 mm of the interface.

We model the virtual quadrant assay on a Cartesian coordinate system whose origin is at the center of the arena, at the intersection between the four quadrants. The concentration field $C(x, y)$ ranges between zero and a maximum, $C_\mathrm{H}$. It is modeled using a hyperbolic tangent function

$C(x,y) = \frac{C_\mathrm{H}}{2}\left[1 + \mathrm{sgn}(x \cdot y)\tanh\left(\frac{d_\mathrm{ax}(x,y)}{5B_\mathrm{w}}\right)\right]$,

which results in a realistic transition and close to zero salt gradients within the quadrants (**Fig. M5**). Here $d_\mathrm{ax}(x, y)$ denotes the distance from the nearest quadrant interface, defined as $\min(|x|, |y|)$ with interfaces at $x = 0$ and at $y = 0$ and $\mathrm{sgn}$ is the sign function.

Choice assay experiments were performed with both naive animals and conditioned animals. In these experiments, animals were washed in either salt-free or salt-containing CTX buffer for 15 minutes before being placed on the assay plate. To mimic pre-exposure in our model, we simulated animals on a plate with a uniform concentration of salt for 15 minutes, and then virtually transferred them to the center of the quadrant plate; when the concentration decrease is encountered, we manually switched on (i.e., recruited) ASH.

As in the experimental protocol, the result was quantified using a chemotaxis index (CI) by counting the number of worms after 600 seconds in each quadrant. In the experiments, worms that remained trapped near a border between quadrants were excluded, and not counted towards the CI. As this does not apply to the model, we did not exclude any region of the plate. To ensure that this does not alter our results, we verified that excluding a region around the border produces statistically identical CIs. Parameter values of the choice assay are given in **Table M5**.

**Table M5.** Virtual choice-assay variable initialization and simulation parameters (naïve animals).

| Variable/Parameter | Value | Description |
|---|---|---|
| N | 1,000 | Number of worms simulated |
| $\tau_{\mathrm{end}}$ | 600 s | Duration of the assay |
| $\mathrm{d}t$ | 0.1 ms | Integration time step of the simulation |
| $\theta(0)$ | Random | Initial direction drawn from uniform distribution |
| $C_{\mathrm{H}}$ | 100 mM | NaCl concentration in NaCl quadrants |
| $B_{\mathrm{w}}$ | 0.2 cm | Half width of boundary region with NaCl gradient |
| $V_i$ | 0 | Initial activity for all neurons |
| $F_i$ | 0 | Initial value for fast component of all sensory neurons |
| $S_i$ | 0 | Initial value for slow component of all sensory neurons |
| $C_0$ | 0 mM | Initial value for the baseline of ASEL |
| $D_{\mathrm{ASER}}$ | 0 | Initial value for fast component in ASER |
| $C_{\mathrm{wash}}$ | 100 mM | Concentration of NaCl in buffer used for conditioning |

**Population models and simulations**

Population models were used to test the robustness of our models and the effect of adaptation on robustness in the quadrant assay. Each population consisted of model animals either with (test) or without (control) ASE adaptation. To generate population models of animals, we perturbed the parameters, $\alpha_{\mathrm{ASEL}}$, $\alpha_{\mathrm{ASER}}$, $\beta_{\mathrm{ASEL}}$, $\beta_{\mathrm{ASER}}$ and $\gamma_{\mathrm{ASEL}}$, $\gamma_{\mathrm{ASER}}$, associated with ASEL and ASER responses (6 parameters in total), by independently subjecting each parameter in each animal to noise. Specifically, model parameters were perturbed by applying multiplicative noise with noise levels independently sampled from a lognormal distribution. In other words, we substituted each parameter $p_{\mathrm{i}}$ by $p_{\mathrm{i}}\ \mathrm{e}^{\sigma x}$ where $x$ was sampled from a normal distribution with standard deviation $\sigma$ (see **Supplementary Fig. 7a**). To focus on the role of adaptation, in these simulations ASH was inactive. We simulated each model animal across each population (with $n = 500$) and compared the CI achieved after 10 minutes of simulation of the quadrant assay for different populations of animals. Noise amplitudes varied from low ($\sigma$ =0.01) to very high noise levels ($\sigma$ =1.28), in which the CI performance falls sharply in both control and test animals.

#### The Model Spot Assay

Simulations were performed in a virtually infinite assay with NaCl spots of standard normal (Gaussian) concentration profiles (**Fig. M6**). We quantified the exploration behavior in terms of hops from one spot to another. For this purpose, the region within standard deviation to a center of a spot is considered “inside” the spot (**Fig. M6**). For each trajectory in the spot assay, we have collected the times the trajectory enters the inside of a spot for the first time after leaving another spot. Supplementary movies S15-S30 show 100 (out of 500) simulated worms, wild-type, ASH-virtual ablated, or no (de) sensitization in ASEL or ASER, on the spot assay, with a peak concentration of 100 or 200 mM NaCl and spot separation distance of 3.3 or 5 cm.

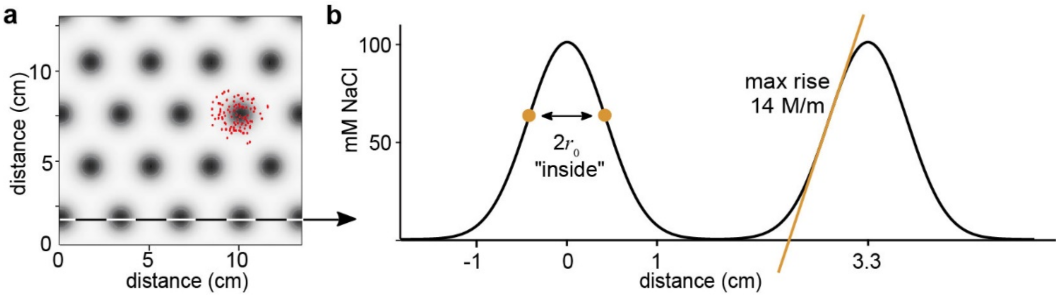


**Fig. M6. The model spot assay.** (**a**) A plate of virtually infinite size is prepared with NaCl spots (in black) arranged in a hexagonal grid. Animals are indicated as red dots. (**b**) The NaCl concentration at a point $p = (x, y)$ is the sum of contributions $\sum_{\text{spots},i} s\,(|p - q_i|)$ from salt spots, with each salt spot following a Gaussian radial concentration profile around a center, denoted $q_i = (x_i, y_i)$, $s(r) = C_\text{S}\exp(-2(r/r_0)^2)$ (assuming radial diffusion is well approximated by linear diffusion from a point). An infinite virtual assay is constructed by making use of the periodicity of the spot positions. At any point, $p$, we approximate the salt concentration by a superposition of the contributions from the seven nearest spots (verified to yield values within machine precision of the exact concentration value).

**Table M6.** Virtual spot assay parameters.

| Initialization (naïve) | Value | Description |
|---|---|---|
| $\tau_\text{end}$ | 3600 s | Duration of the assay |
| $\text{d}t$ | 0.1 ms | Integration time step of the simulation |
| $\theta(0)$ | Random | Initial direction (sampled from a uniform distribution) |
| $C_\text{S}$ | 100 mM | Peak NaCl concentration (at each spot center) |
| $r_0$ | 0.6 cm | Spot radius ([NaCl] standard deviation) |

**Supplementary Movies**

Movies S1-S14 show 500 simulated worms for 10 minutes on the quadrant choice assay, with upper right and lower left quadrants containing 100 mM NaCl, and the other two quadrants contain no NaCl. Corresponds to Figure 8.

Movie S1: ASH ablated
Movie S2: ASH ablated, no (de)sensitization in ASEL or ASER
Movie S3: ASH ablated, no connection between ASEL and steering
Movie S4: ASH ablated, no connection between ASEL and pirouettes
Movie S5: ASH ablated, no connection between ASER and steering
Movie S6: ASH ablated, no connection between ASER and pirouettes
Movie S7: ASH ablated, ASEL ablated
Movie S8: ASH ablated, ASER ablated
Movie S9: ASH ablated, no steering
Movie S10: ASH ablated, ASEL ablated, no steering
Movie S11: ASH ablated, ASER ablated, no steering
Movie S12: ASH ablated, no pirouettes
Movie S13: ASH ablated, ASEL ablated, no pirouettes
Movie S14: ASH ablated, ASER ablated, no pirouettes

Movies S15-S18 show 100 (out of 500) simulated worms on the spot assay, with a peak concentration of 100 mM NaCl and spot separation distance of 3.3 cm.

Movie S15: wild type
Movie S16: ASH ablated
Movie S17: no (de)sensitization in ASEL or ASER.
Movie S18: ASH ablated, with no (de)sensitization in ASEL or ASER.

Movies S19-S22 show 100 simulated worms on the spot assay, with a peak concentration 200 mM and spot separation distance of 3.3 cm.

Movie S19: wild type simulation
Movie S20: ASH-virtual ablated simulations.
Movie S21: no (de)sensitization in ASEL or ASER.
Movie S22: ASH-virtual ablated simulations, with no (de)sensitization in ASEL or ASER.

Movies S23-S26 show 100 simulated worms on the spot assay, with a peak concentration100 mM and spot separation distance of 5.0 cm.

Movie S23: wild type simulation
Movie S24: ASH-virtual ablated simulations.
Movie S25: no (de)sensitization in ASEL or ASER.
Movie S26: ASH-virtual ablated simulations, with no (de)sensitization in ASEL or ASER.

Movies S27-S30 show 100 simulated worms on the spot assay, with a peak concentration 200 mM and spot separation distance of 5.0 cm.

Movie S27: wild type simulation

Movie S28: ASH-virtual ablated simulations.

Movie S29: no (de)sensitization in ASEL or ASER.

Movie S30: ASH-virtual ablated simulations, with no (de)sensitization in ASEL or ASER.
